# A Novel Support Vector Machine-Based One-Day, Single-Dose Prediction Model of Genotoxic Hepatocarcinogenicity in Rats

**DOI:** 10.1101/2024.02.19.579908

**Authors:** Min Gi, Shugo Suzuki, Masayuki Kanki, Masanao Yokohira, Tetsuya Tsukamoto, Masaki Fujioka, Arpamas Vachiraarunwong, Guiyu Qiu, Runjie Guo, Hideki Wanibuchi

**Affiliations:** Department of Environmental Risk Assessment, Graduate School of Medicine, Osaka Metropolitan University, Osaka 545-8585, Japan; Department of Molecular Pathology, Graduate School of Medicine, Osaka Metropolitan University, Osaka 545-8585, Japan; Department of Medical Education, Faculty of Medicine, Kagawa University, Miki-cho, Kita-gun, Kagawa 761-0793, Japan; Department of Pathology and Host-Defense, Faculty of Medicine, Kagawa University, Miki-cho, Kita-gun, Kagawa 761-0793, Japan; Department of Diagnostic Pathology, Graduate School of Medicine, Fujita Health University, Toyoake, Aichi 470-1192, Japan

**Author notes:** **Corresponding author** Hideki Wanibuchi.

**Keywords:** Genotoxic hepatocarcinogen, genotoxicity and carcinogenicity prediction, SVM-based predictive classifier, a single-dose, one-day rat model, prediction marker gene

## Abstract

The development of a rapid and accurate model for determining the genotoxicity and carcinogenicity of chemicals is crucial for effective cancer risk assessment, and it also contributes to cancer prevention. This study aims to develop a one-day, single-dose model for identifying genotoxic hepatocarcinogens (GHCs) in rats. Microarray gene expression data from the livers of rats administered a single dose of 58 compounds, including 5 GHCs, was obtained from the Open TG-GATEs database and used for the identification of marker genes and the construction of a predictive classifier to identify GHCs in rats. We identified 10 gene markers commonly responsive to all 5 GHCs and used them to construct a support vector machine-based predictive classifier. This classifier effectively distinguishes GHCs from other compounds, demonstrating 100% sensitivity and over 96% specificity. To further assess the model’s effectiveness and reliability, we conducted multi-institutional one-day single oral administration studies on rats. These studies examined 64 compounds, including 23 GHCs, with gene expression data of the marker genes obtained via quantitative PCR (qPCR) 24 hours after a single oral administration. Our results demonstrate that qPCR analysis is an effective alternative to microarray analysis. The GHC predictive model showed high accuracy and reliability, achieving a sensitivity of 91% (21/23) and a specificity of 93% (38/41) across multiple validation studies in three institutions. In conclusion, the present one-day single oral administration model proves to be a reliable and highly sensitive tool for identifying GHCs and is anticipated to be a valuable tool in identifying and screening potential GHCs.

## Introduction

Evaluation of genotoxicity and carcinogenicity is an essential part of the developmental process for pharmaceuticals, food additives, agricultural chemicals, and industrial chemicals. Various experimental models have been established for assessing the genotoxicity and carcinogenicity of chemicals. The 2-year rodent carcinogenicity bioassay has historically been the gold standard for predicting the carcinogenic hazard of chemicals to humans. However, 2-year studies require preliminary dose range finding studies and the overall time required to complete a 2-year study is generally 3-4 years, and the studies require significant quantities of the test material. These factors make 2-year studies unsuitable for the evaluation of materials during early development stages. Two-year rodent studies are also very costly, labor-intensive, and time-consuming and require a large number of animals. This severely limits the ability to carry out large-scale evaluation of the carcinogenicity of multiple chemical compounds using 2-year studies. In addition, the relevance and necessity of conducting of 2-year carcinogenicity studies has been debated (Cohen 2010b; Ennever and Lave 2003). However, alternative testing methods, such as middle-term carcinogenicity tests (Ashby 1996; Cohen 2010a; Cohen et al. 2001; Ito et al. 2003), often require special procedures or the use of genetically modified animals, which limits the number of facilities capable of conducting these tests (Eastmond et al. 2013).

Genotoxicity is increasingly being recognized as a bona fide toxicological endpoint (Hsieh et al. 2019) and is a crucial factor in extrapolating carcinogenic risk from experimental animals to humans based on the mode of action, particularly for estimating risks associated with low-dose exposures (Cohen et al. 2019; Nohmi 2018; O’Brien et al. 2006; SCHER 2009). While threshold extrapolation is accepted for non-genotoxic carcinogens, as they do not exhibit carcinogenic effects at low doses, linear non-threshold extrapolation is generally accepted for genotoxic carcinogens, suggesting some risk at any exposure level (EPA 2005). However, recent trends propose introducing a threshold concept for genotoxic carcinogens, especially those causing indirect genetic toxicity through mechanisms like chromosomal abnormalities, oxidative stress, and DNA synthesis inhibition (Hartwig et al. 2020; SCHER 2009). Consequently, it is generally accepted that direct genotoxic carcinogens should not be used as food additives, pesticides, or veterinary drugs, or used for pharmaceuticals, except for anti-cancer drugs (Bolt 2008; Nohmi 2018; WHO and FAO 2021).

Well-established genotoxicity assays include the bacterial reverse mutation test, in vitro and in vivo tests that measure chromosomal aberrations, and micronucleus formation (Beal et al. 2023; Kirkland et al. 2005). While these assays are high throughput, they lack organ specificity. *In vivo* transgenic rodent models are capable of determining organ-specific mutagenesis induced by carcinogens (Nohmi et al. 2000), but these methods are laborious, time-consuming, and expensive. Currently, using multiple genotoxicity tests to assess the genotoxicity of chemicals is generally adopted as no single test is capable of detecting all genotoxic mechanisms relevant to tumorigenesis (Beal et al. 2023; ICH 2008; Kirkland et al. 2007).

Given the annual production of tens of thousands of new chemical compounds, establishing a comprehensive system that rapidly and accurately verifies both genotoxicity and carcinogenicity of chemicals is crucial. Gene expression data has proven to be valuable in predicting the carcinogenicity of chemicals (Corton et al. 2022; Ellinger-Ziegelbauer et al. 2009; Furihata et al. 2016; Gant et al. 2023; Waters et al. 2010). Recent technological advancements have enabled the development of machine learning algorithms, such as Support Vector Machine (SVM), Random Forest, and Decision Tree, for predicting carcinogenicity using gene expression data.

The liver, being the most frequently targeted site for chemical carcinogens (Gold et al. 2001; Gold et al. 1991), has been the focus of many successful hepatocarcinogenicity prediction models using toxicogenomics data in rats (Corton et al. 2020; Ellinger-Ziegelbauer et al. 2009; Furihata et al. 2016; Hill et al. 2020; Uehara et al. 2011). These models are particularly suitable at evaluating hepatocarcinogenicity and predicting non-genotoxic hepatocarcinogens (NGHC) (Ellinger-Ziegelbauer et al. 2008; Fielden et al. 2011; Kanki et al. 2016; Smith et al. 2020; Uehara et al. 2011; Yamada et al. 2016). However, the validation of these models for predicting genotoxic hepatocarcinogens (GHC) remains insufficient, partly due to the small number of GHCs tested (Ellinger-Ziegelbauer et al. 2008; Smith et al. 2020; Uehara et al. 2011; Waters et al. 2010). Consequently, a reliable and practical prediction model for GHCs has not yet been reported in the published literature. Consequently, current hepatocarcinogenicity prediction models require improvement. Current models require a large number of marker genes, and this necessitates the use of microarray analysis. However, microarray analyses leads to high costs and difficulties in validation at different institutions, limiting the number of compounds that can be tested. Addressing these challenges is crucial for advancing the development of effective and efficient GHC prediction models.

The aim of the present study is to develop a one-day, single-dose model for predicting genotoxic hepatocarcinogenicity of test compounds in rats, utilizing gene expression data obtained via quantitative PCR (qPCR) 24 hours after a single oral administration of the test material. Initially, we identified a set of 10 gene markers to predict the hepatocarcinogenicity of GHCs and constructed a prediction model using the SVM algorithm. This model was validated with gene expression data derived from microarray analysis of rat livers treated with 58 compounds, including 5 known GHCs, sourced from the Open Toxicogenomics Project-Genomics Assisted Toxicity Evaluation System (Open TG-GATEs) database. To assess the predictive accuracy of our model, and with the goal of reducing costs and analysis time, and improving reproducibility at different institutions, we conducted multi-institutional one-day single oral dose studies with rats. These studies examined 64 compounds, including 23 GHCs, analyzing the expression data of the marker genes using qPCR.

## Materials and methods

### 1. Classification of test compounds

Compounds were classified into two categories based on their mutagenicity and hepatocarcinogenicity in rats: GHCs and other compounds. Compounds that tested positive in the Ames test and were carcinogenic to the livers of rats were defined as GHCs. The OTHER category included a variety of compounds, encompassing nongenotoxic hepatocarcinogens, nongenotoxic nonhepatocarcinogens (NGNHC) that are carcinogenic to organs other than the liver, genotoxic nonhepatocarcinogens (GNHC) that are carcinogenic to organs other than the liver, genotoxic noncarcinogens (GNC), and nongenotoxic noncarcinogens (NGNC). Information regarding the CAS numbers and suppliers of the chemicals tested in the validation studies is provided in Supplementary Table 1.

Information regarding the carcinogenicity and mutagenicity of these compounds was obtained from the National Toxicology Program (https://ntp.niehs.nih.gov) and the ToxInfo.io databases (https://www.toxinfo.io). ToxInfo.io is a comprehensive toxicology and hazardous chemicals database, with multiple integrated databases including the Carcinogenic Potency Database (CPDB), Chemical Carcinogenesis Research Information System (CCRIS), Integrated Risk Information System, and Genetic Toxicology Data Bank (GENE-TOX).

### 2. Gene expression data for identification of marker genes and construction of a GHC classifier

Microarray gene expression data from the livers of Sprague-Dawley rats exposed to 58 compounds were utilized for the identification of marker genes and the construction of a classifier to predict GHC in rats. This data was obtained from the Open TG-GATEs, a public toxicogenomic database developed in Japan (Igarashi et al. 2015).

In the Open TG-GATEs database, each compound was evaluated at three dose levels along with a concurrent control in single oral administration studies. In these studies, rats were administered a compound and then sacrificed at 3, 6, 9, or 24 hours post-administration. The highest dose level used was the maximum tolerated dose that was identified in a dosing study. The three dose levels were multiples of the square root of 10.

### 3. Identification of marker genes for a predictive classifier

To identify marker genes capable of distinguishing GHC in rats, we analyzed microarray gene expression data of five GHCs deposited in the Open TG-GATEs database: 2-Acetamidofluorene (AAF), N-Nitrosodiethylamine (NDEA), 2-Nitrofluorene (2-NF), Aflatoxin B1 (ATB), and N-Nitrosomorpholine (NMP) (Table 1). The low doses of these GHCs used in the 24-hour single oral administration toxicity studies were significantly higher – 82, 377, 351, 313, and 917 times, respectively – than their TD50 values reported in the Carcinogenic Potency Database (CPDB). Our analysis was concentrated on identifying common differentially expressed genes in response to these doses.

**Table 1.** TD50 and Doses of GHCs used in single dose toxicity studies documented in the OPEN TG-GATEs database.

| GHC | TD50 (mg/kg bw) <sup>a</sup> | Doses (mg/kg bw) <sup>b</sup> |  |  |
| --- | --- | --- | --- | --- |
|  |  | Low | Middle | High |
| 2-Acetamidofluorene (2-AAF) | 1.22 | 100* (82 <sup>c</sup>) | 300 (246) | 1000 (820) |
| N-Nitrosodiethylamine (NDEA) | 0.0265 | 10* (377) | 30 (1132) | 100 (3774) |
| 2-Nitrofluorene (2-NF) | 0.285 | 100* (351) | 300 (1053) | 1000 (3509) |
| Aflatoxin B1 (AFB1) | 0.0032 | 1* (313) | 3 (938) | 10 (5938) |
| N-nitrosomorpholine (NMP) | 0.109 | 100* (917) | 300 (2752) | 600 (5505) |

Abbreviation: GHC: genotoxic hepatocarcinogen.
<sup>a</sup> TD50 refers to the daily dose in mg/kg/bw/day administered for life to induce tumors in half of the test animals that would have remained tumor-free at zero dose: Reported in the Carcinogenic Potency Data Base (CPDB).
<sup>b</sup> Doses used in the single dose toxicity studies in the OPEN TG-GATEs.
<sup>c</sup> Values in parentheses represent multiples of the dose relative to the TD50.
\* Marker genes were identified as those that consistently showed differential expression in response to low doses of all the five GHCs.

The methodology for analyzing the microarray gene expression data was as described previously (Kanki et al. 2016). In brief, three CEL files per dosing group, generated with an Affymetrix Rat Genome 230 2.0 array (Affymetrix, Santa Clara, CA, USA), were obtained from the Open TG-GATEs. The Microarray Analysis Suite 5.0 (MAS; Affymetrix) was employed to quantify each probe set signal. Raw signal data was normalized using the median value of all probe sets (per chip normalization). GeneSpring GX software (Version 12.6; Agilent Technologies, Inc., Santa Clara, CA, USA) was utilized for normalization and statistical analysis. Differentially expressed genes were identified using Welch’s t-test (p<0.05) and a criterion of 2-fold change in expression compared to controls.

### 4. Ingenuity pathway analysis

The gene network analysis for the selected 10 marker genes was conducted using Ingenuity Pathway Analysis Software (Version 01-22-01; Ingenuity Systems, Qiagen, CA, USA). This analysis aligned the marker genes with established genetic pathways and networks, aiming to illustrate their potential biological roles and interactions relevant to genotoxic hepatocarcinogenicity.

### 5. Construction of a GHC classifier using microarray gene expression data from the Open TG-GATEs

A predictive classifier for GHC was constructed using the SVM algorithm, a type of supervised machine learning algorithm. This was integrated into the GeneSpring GX software. In the SVM analysis, we applied a linear kernel type and used leave-one-out cross-validation. The expression data of the 10 marker genes, obtained from the Open TG-GATEs database, were used for both constructing and testing the classifier. A compound predicted as a GHC by the classifier was deemed positive, while those predicted as OTHER were deemed negative.

To construct the predictive classifier, training was conducted using known GHC and OTHER compounds. For the positive training dataset, we selected microarray expression data of the marker genes at low doses of 3 GHCs: 2-AAF, NDEA, and 2-NF. For the negative training dataset, we used microarray expression data of marker genes at low doses of 14 OTHER compounds: 3 NGHCs - Thioacetamide (TAA), Hexachlorobenzene (HCB), and Wy-14643 (WY); 2 GNHCs - Cyclophosphamide (CPA) and Nitrofurantoin (NFT); 3 NGNHCs - Butylated hydroxyanisole (BHA), Methimazole (MTZ), and Sulfasalazine (SS); and 6 NGNCs - Allyl Alcohol (AA), Cyclosporine A (CA), Furosemide (FUR), Promethazine (PMZ), Sulindac (SUL), and Tetracycline (TC).

### 6. In silico validation of the GHC predictive classifier using microarray gene expression data from the Open TG-GATEs

To validate the GHC predictive classifier, we used marker gene expression data for 5 GHCs and 53 OTHER compounds from the Open TG-GATEs database (Table 2).

**Table 2.**
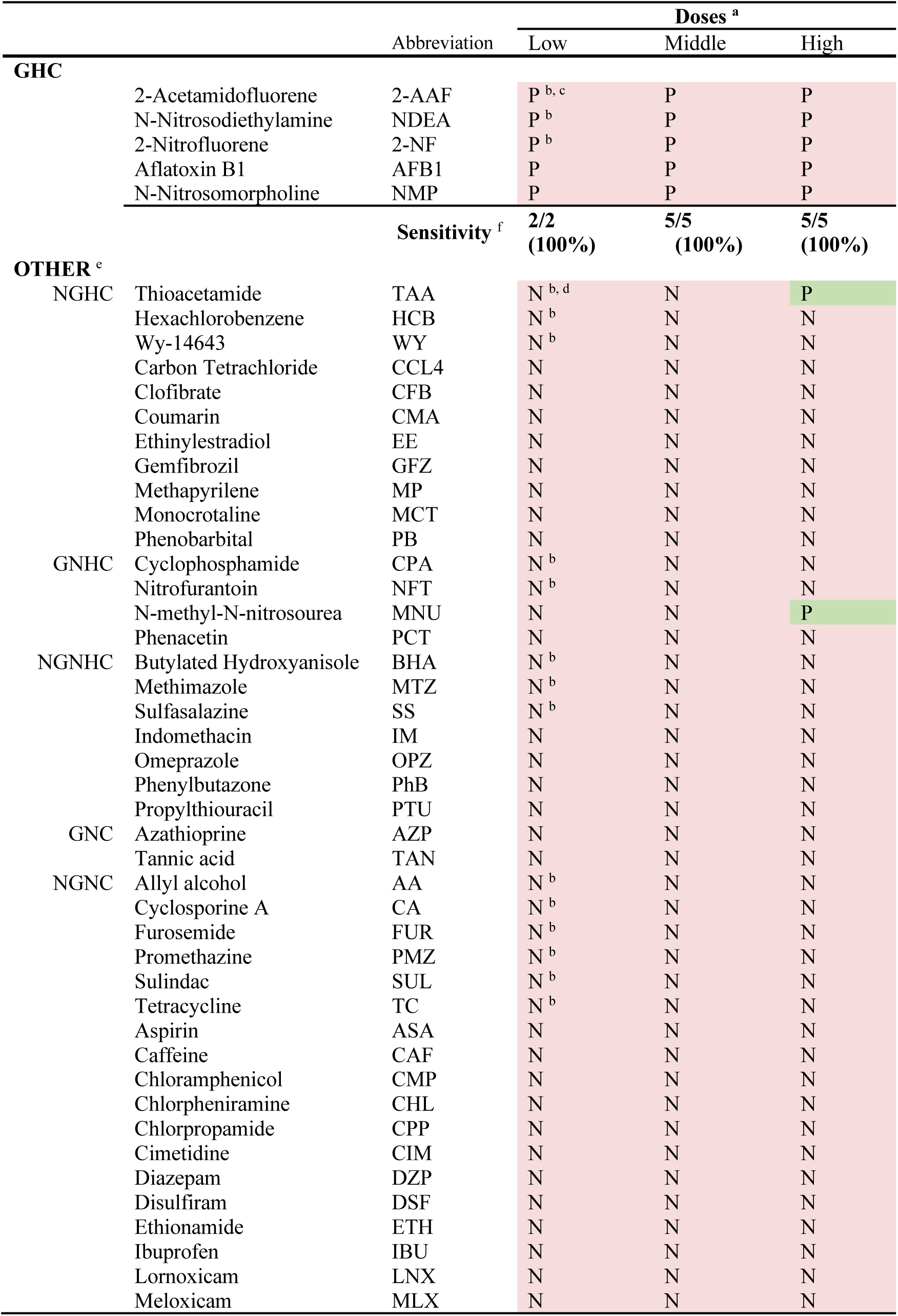
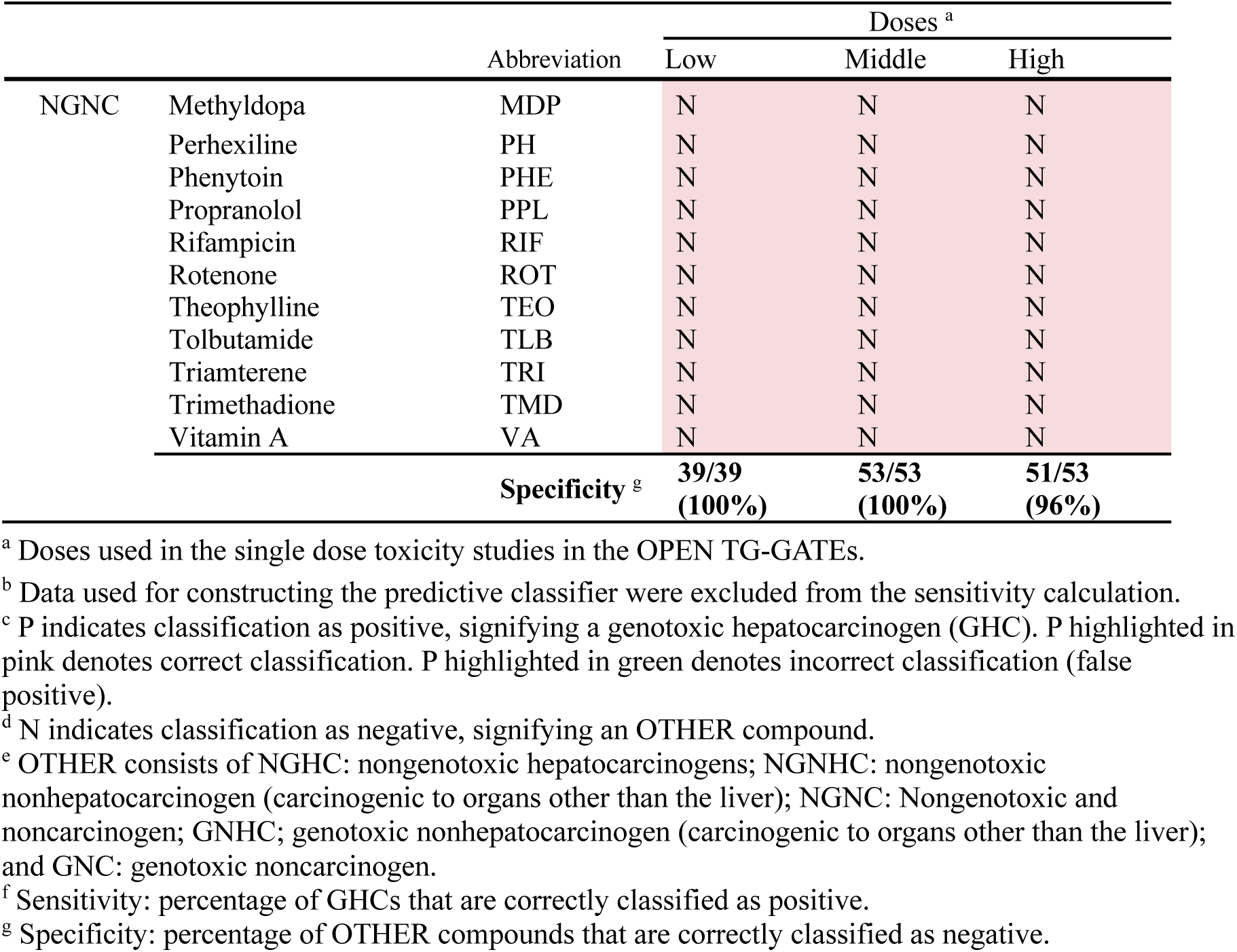
Results of the GHC predictive classifier using microarray gene expression data of 58 compounds obtained from Open TG-GATEs.

For the positive test set (GHC dataset), we used marker gene expression data from middle and high doses of 2-AAF, NDEA, and 2-NF, and data from low, middle, and high doses of AFB1 and NMP.

For the negative test set (OTHER compounds dataset), marker gene expression data from middle and high doses of the 14 OTHER compounds used in the construction of the GHC predictive classifier were used. In addition, data from low, middle, and high doses of 39 OTHER compounds were used: 8 NGHCs - Carbon Tetrachloride (CCL4), Clofibrate (CFB), Coumarin (CMA), Ethinylestradiol (EE), Gemfibrozil (GFZ), Methapyrilene (MP), Monocine (MCT), and Phenobarbital (PB); 2 GNHCs - N-methyl-N-nitrosourea (MNU) and Phenacetin (PCT); 4 NGNHCs - Indomethacin (IM), Omeprazole (OPZ), Phenylbutazone (PhB), and Propylthiouracil (PTU); 2 GNCs - Azathioprine (AZP) and Tannic Acid (TAN); 23 NGNCs - Aspirin (ASA), Caffeine (CAF), Chloramphenicol (CMP), Chlorpheniramine (CHL), Chlorpropamide (CPP), Cimetidine (CIM), Diazepam (DZP), Disulfiram (DSF), Ethionamide (ETH), Ibuprofen (IBU), Lornoxicam (LNX), Meloxicam (MLX), Methyldopa (MDP), Perhexiline (PH), Phenytoin (PHE), Propranolol (PPL), Rifampicin (RIF), Rotenone (ROT), Theophylline (TEO) (Mateo et al.), Tolbutamide (TLB), Triamterene (TRI), Trimethadione (TMD) and Vitamin A (VA).

For the in silico validation of the constructed GHC predictive classifier, the sensitivity and specificity of the predicted compounds were calculated. Prediction results of training data at the low doses were not included in the calculation of the sensitivity and specificity. For the middle and high doses, all of the prediction results from above compounds (5 GHCs and 53 OTHER compounds) were applied to calculate the sensitivity and specificity.

### 7. The first set of validation studies (Validation Study 1): Evaluation of 5 GHCs in one-day single dose oral administration studies

#### 7-1 Test compounds

To validate the GHC predictive classifier and assess whether the marker gene expression data obtained via qPCR could be an alternative to microarray analysis data, we selected 5 GHCs not previously evaluated in the Open TG-GATEs for validation. These GHCs were 2-Aminoanthraquinone (2-AA), 2-Nitropropane (2-NP), 2,4-Diaminotoluene (2,4-DAT), 2-Amino-3,8-dimethylimidazo[4,5-*f*]quinoxaline (MeIQx), and 2-amino-3-methylimidazo[4,5-*f*]quinoline (Table 3).

**Table 3.** Doses and prediction results of Validation Study 1.

|  |  | Doses |  |  |  |  |
| --- | --- | --- | --- | --- | --- | --- |
|  |  | Low<br>(1/30 of LD50) | Middle<br>(1/10 of LD50) | High<br>(1/3 of LD50) |  | LD50 <sup>a</sup> |
|  |  | qPCR | qPCR | qPCR | Microarray analysis | qPCR |
| 2-Aminoanthraquinone | 2-AA <sup>b</sup> | N <sup>d</sup> | N | N | N | P |
| 2-Nitropropane | 2-NP <sup>b</sup> | N | P <sup>e</sup> | P | P | NE <sup>f</sup> |
| 2,4-Diaminotoluene | 2,4-DAT <sup>b</sup> | N | N | P | P | NE |
| 2-Amino-3,8-dimethylimidazo[4,5-f]quinoxaline | MeIQx <sup>c</sup> | N | P | P | P | NE |
| 2-Amino-3-methylimidazo[4,5-f]quinoline | IQ <sup>c</sup> | N | P | P | P | NE |

<sup>a</sup> Lethal dose 50 (LD50) values for oral administration in rats are 1500 mg/kg bw for 2-AA, 720 mg/kg bw for 2-NP, and 135 mg/kg bw for 2,4-DAT. LD50 values MeIQx and IQ are not available.
<sup>b</sup> Low, middle, and high doses for 2-AA, 2-NP, and 2,4-DAT were set at approximately 1/30; 1/10, and 1/3 of their respective LD50 values. The doses were as follows: 2-AA: 50, 150, and 450 mg/kg bw; 2-NP: 24, 72, and 240 mg/kg bw; 2,4-DAT: 4.5, 13.5, and 45 mg/kg bw.
<sup>c</sup> Middle dose for MeIQx and IQ was set at 10 times of their respective carcinogenic doses. The low and high doses were determined as 1/3 and 3 times the middle dose. The doses were as follows: MeIQx: 50, 160, and 480 mg/kg bw; IQ: 40, 120, and 360 mg/kg bw.
<sup>d</sup> N indicates classification as negative, signifying an OTHER compound. N highlighted in green denotes incorrect classification (false negative).
<sup>e</sup> P indicates classification as positive, signifying a genotoxic hepatocarcinogen (GHC). P highlighted in pink denotes correct classification.
<sup>f</sup> NE: not examined.

#### 7-2 Animals

Animal experiments were conducted in Osaka Metropolitan University (referred to as Institution O). Five-week-old male SD rats were obtained from Charles River Laboratories Japan, Inc. (Hino, Siga, Japan). The rats were housed in polycarbonate cages (5 rats per cage) with wood chips for bedding. The experimental animal room was maintained at a temperature of 22 ± 3°C, relative humidity of 55 ± 5%, and a 12-h light/dark cycle. The rats underwent a 7-day acclimatization period and were 6 weeks old at the beginning of the study.

#### 7-3 Experimental protocol

The experimental protocol was reviewed and approved by the Ethics Review Committee for Animal Experimentation and conducted in accordance with the Guidelines for Proper Conduct of Animal Experiments (Science Council of Japan, 2006). CE2 basal pellet diet (Clea Japan, Inc., Tokyo, Japan.) and tap water were available *ad libitum* throughout the study.

Groups of 5 rats were administered test compounds or vehicle (for the control group) by oral gavage. The test compounds were dissolved or suspended in 0.5% methylcellulose (vehicle) at target doses with an administration volume of 5 ml/kg bw. Prior to administration, the body weight of each rat was measured.

In the first experiment of Validation Study 1, doses for 2-AA, 2-NP, and 2,4-DAT were determined based their oral acute toxicity in rats. Doses were set at 1/30; 1/10, and 1/3 of the oral lethal dose (LD50). Doses were based on the assumption that the desired dose for the 24-hour single dose administration study is the maximum tolerated dose. Oral LD50 values for these GHCs were obtained from https://pubchem.ncbi.nlm.nih.gov/. For MeIQx and IQ, whose oral LD50 values were not available, the middle dose was set at 10 times their respective carcinogenic doses (NTP 2002; Takayama et al. 1984). Subsequently, the low and high doses were set at 1/3 and 3 times the middle dose. Vehicle-treated groups were included as a negative control.

In the second experiment of Validation Study 1, the LD50 dose of 2-AA was evaluated because in the first experiment it was predicted as negative at doses up to 1/3 of the LD50, and no apparent toxicities were observed. Vehicle-treated groups were included as a negative control.

Twenty-four hours after treatment, rats were euthanized via transection of the abdominal aorta under deep anesthesia induced by inhalation of an overdose of isoflurane (Abbott Japan Co., LTD., Tokyo, Japan), using a Small Animal Anesthetizer (MK-A110D, Muromachi Kikai Co., LTD., Tokyo, Japan) coupled with an Anesthetic Gas Scavenging System (MK-T 100E, Muromachi Kikai Co., LTD., Tokyo, Japan). The animals were not fasted before euthanasia. At necropsy, fresh liver tissue from the left lateral lobe was submerged in RNAlater™ Stabilization Solution (Thermo Fisher Scientific Baltics UAB, Vilnius, Lithuania) and stored at 4°C for one day, then stored at −80°C until RNA isolation. Three sections of liver tissue (one each from the left lateral lobe, right middle lobe, and caudate lobe) were fixed in phosphate-buffered formalin, embedded in paraffin, and processed for hematoxylin and eosin staining. The remaining liver tissues were snap frozen in liquid nitrogen and stored at −80°C.

Microarray analysis was conducted on three liver samples each from the control and high-dose groups of each test compound in the first experiment (450 mg/kg for 2-AA; 240 mg/kg for 2-NP; 45 mg/kg for DAT; 480 mg/kg for MeIQx; and 360 mg/kg for IQ) using Affymetrix Rat Genome 230 2.0 arrays (Affymetrix, Santa Clara, CA, USA) as described previously (Kanki et al. 2016). The microarray gene expression data was analyzed as described above, using the Microarray Analysis Suite 5.0 and GeneSpring GX software. In addition, mRNA expression analysis of the 10 marker genes was conducted on all liver samples from both the first and the second experiments using qPCR (Table 3).

#### 7-4 mRNA Expression Analysis in Rat Livers

Total RNA was extracted from the liver tissues preserved in RNAlater using the RNeasy mini kit (QIAGEN) following the manufacturer’s instructions. For qPCR, cDNA synthesis was performed using 1 µg of RNA with an Invitrogen™ SuperScript VI VILO™ Master Mix (Thermo Fisher Scientific Inc., MA, USA). Sequence-specific primers and probes for each gene were procured from Roche Applied Science, Mannheim, Germany (Supplementary Table 2).

Expression assays were conducted in a 20 µl reaction mixture containing 1 µl cDNA, 1.2 µl of a mixture of forward and reverse primers and probe (0.2 µM each), and 10 µl TaqMan Fast Universal PCR Master Mix. These assays were performed using an Applied Biosystems™ 7500 Fast Real-Time PCR System. The Eukaryotic 18S rRNA Endogenous Control (VIC™/MGB probe, primer limited) was used as an internal control (Thermo Fisher Scientific Inc., MA, USA). Expression analysis was performed in triplicate for each sample.

The gene expression values for each sample were calculated using the equation 2^-ΔCt (ΔCt = Ct average marker gene - Ct average 18S). The relative gene expression ratios between treated and control samples were then converted into log base 2 and the average of the log base 2 values in each dose group was applied to the GHC predictive classifier.

### 8. The second set of validation studies (Validation Study 2): Multi-Institution validation studies using marker gene expression data of 60 compounds obtained from qPCR analyses in one-day single dose oral administration studies

#### 8-1 Test compounds

A total of 60 compounds, comprising 19 GHCs and 41 OTHER compounds, were included in Validation Study 2 (Table 4). The 19 GHCs consisted of 2-NP, 3-Methyl-4-dimethylaminoazobenzene (3’-MeDAB), 4,4’-Methylene-bis(2-chloro-aniline) (MOCA), 4,4’-Thiodianiline (TDA), Acid Red 26 (AR-26), Auramine-O (AO), N-Dimetylnitorosamine (DMN), N-Nitrosodiethanolamine (NDEAL), NDEA, N-Dimetylnitorosamine (NDMA), N-Nitrosoethylmethylamine (NEMA), N-nitrosopyrrolidine (NPYR), Nitrosodibutylamine (NDBA), *o*-Aminoazotoluene (AAT),

**Table 4.**
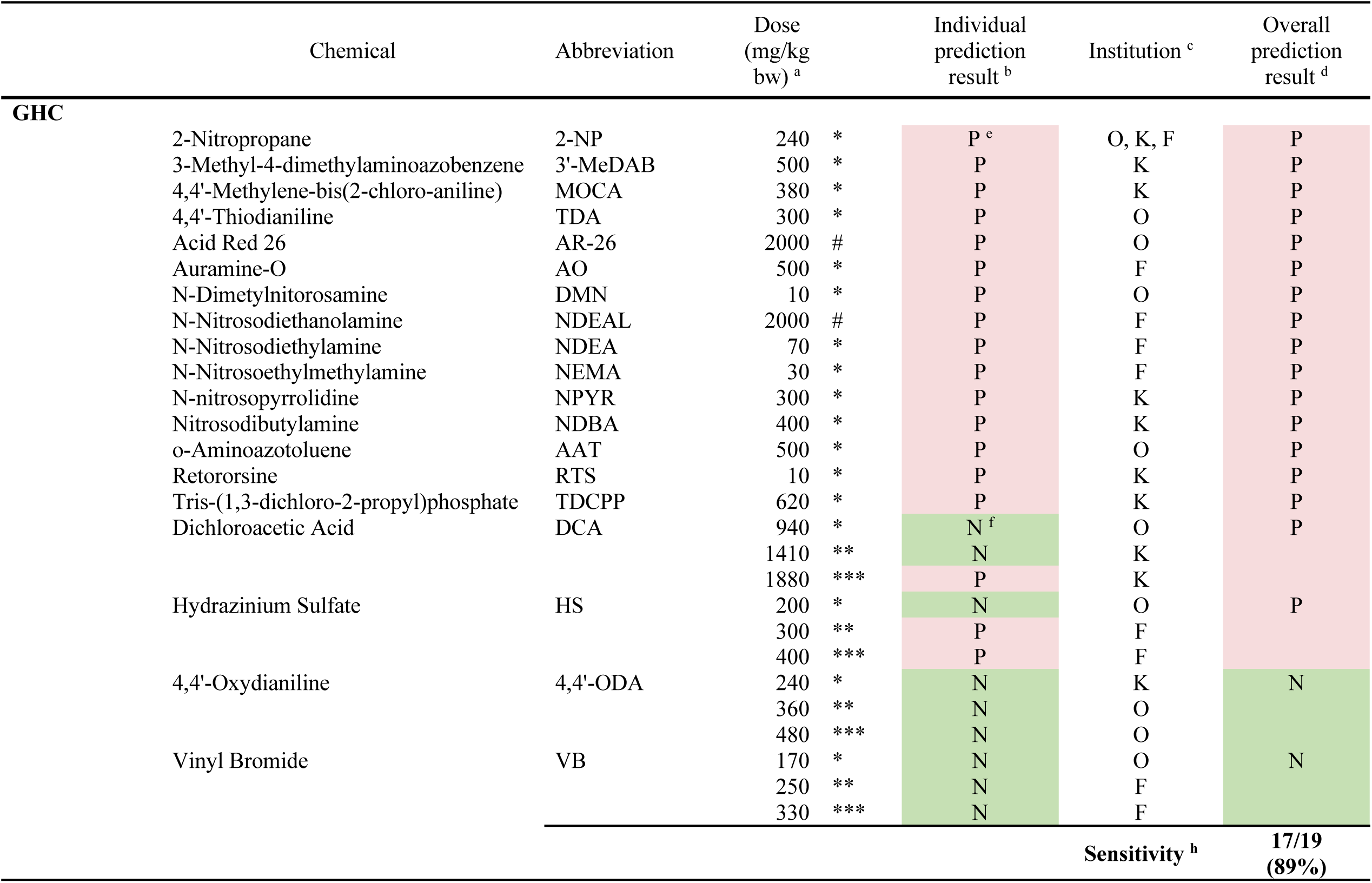
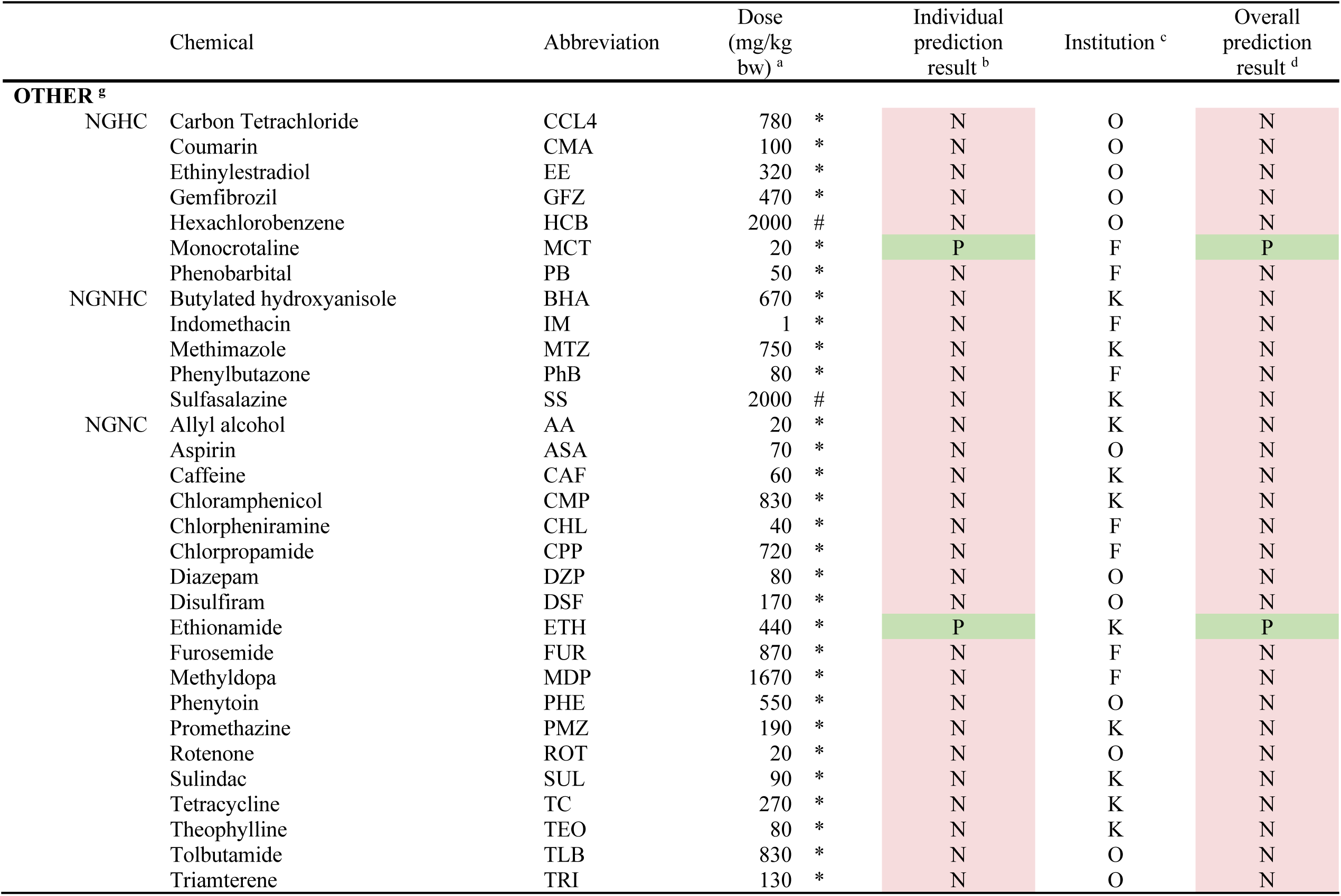
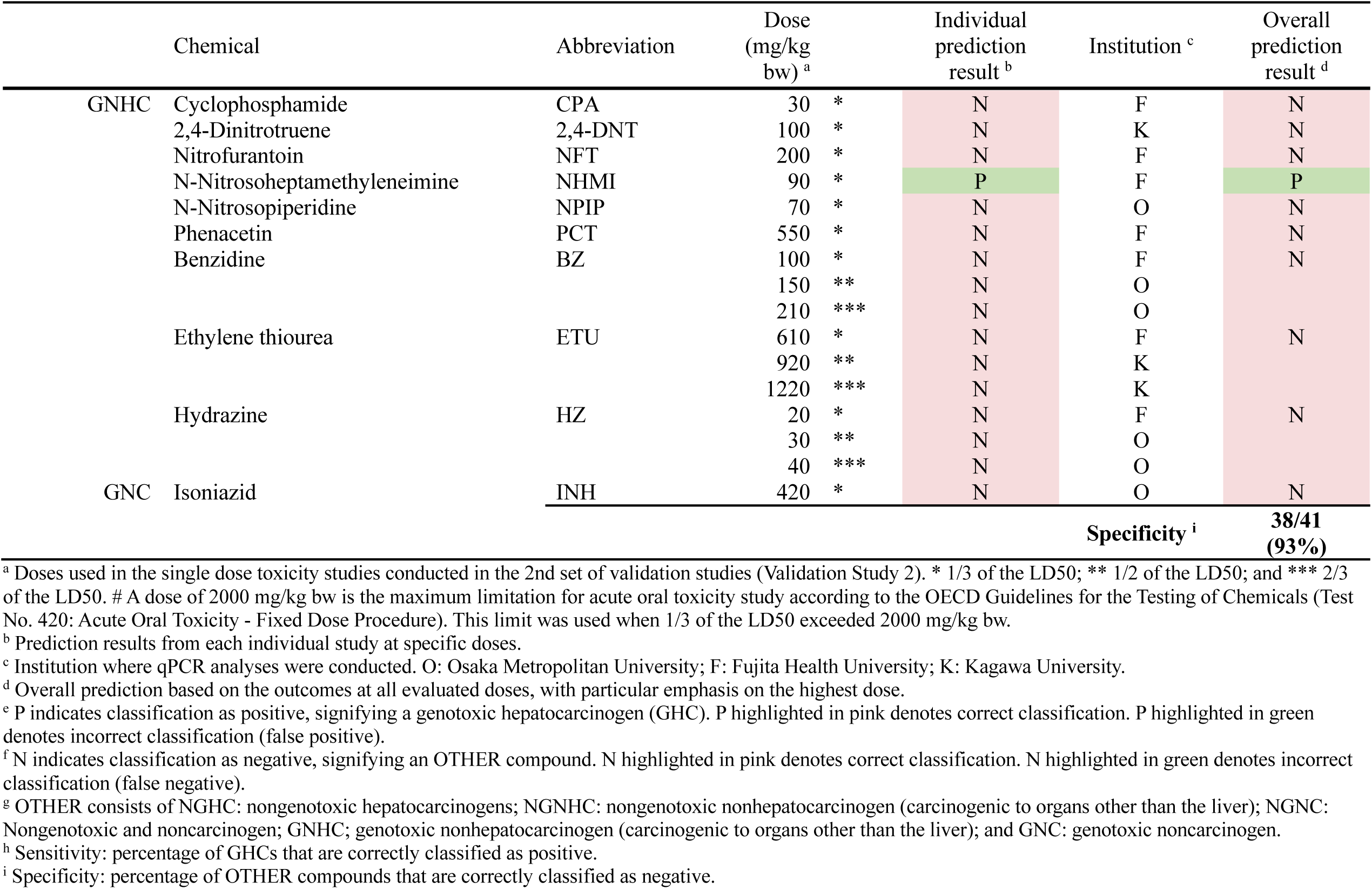
Prediction results of Validation Study 2 using marker gene expression data obtained by qPCR analysis.

Retororsine (RTS), Tris-(1,3-dichloro-2-propyl)phosphate (TDCPP), Dichloroacetic Acid (DCA), Hydrazinium sulfate (HS), 4,4’-Oxydianiline (4,4’-ODA), and Vinyl Bromide (VB).

The 41 OTHER compounds consisted of 7 NGHCs - CCL4, CMA, EE, GFZ, HCB, MCT, and PB; 5 NGNHCs – BHA, IM, MTZ, PhB, and SS; 19 NGNCs – AA, ASA, CAF, CMP, CHL, CPP, DZP, DSF, ETH, FUR, MDP, PHE, PMZ, ROT, SUL, TC, TEO, TLB, and TRI; 9 GNHCs – CPA, 2,4-Dinitrotruene (2,4-DNT), NFT, N-Nitrosoheptamethyleneimine (NHMI), N-Nitrosopiperidine (NPIP), PCT, Benzidine (BZ), Ethylene thiourea (ETU), and Hydrazine (HZ); 1 GNC – Isoniazid (INH).

#### 8-2 Animals

Five-week-old male SD rats were obtained from Charles River Laboratories Japan, Inc. (Hino, Siga, Japan). Rats underwent a 7-day acclimatization period and were 6 weeks old at the beginning of the study.

#### 8-3 Experimental protocol

Animal experiments were conducted at three institutions: Institution O, Kagawa University (referred to as Institution K), and DIMS Medical Science Institute, Inc. (Ichinomiya, Japan): DIMS Medical Science Institute conducted the animal experiments for Fujita Health University (referred to as Institution F). All three institutions followed the same protocols described above for the animal study. The experimental protocol was reviewed and approved by the Ethics Review Committee for Animal Experimentation at each respective institution and conducted in accordance with the Guidelines for Proper Conduct of Animal Experiments as laid out by the Science Council of Japan in 2006.

RNA extraction and gene expression assays were performed following the same protocols described in the Validation Study 1 at three institutions: Institution O, Institution K, and Institution F, using an Applied Biosystems™ 7500 Fast Real-Time PCR System at Institution O, an Applied Biosystems™ StepOnePlus™ Real-Time PCR System at Institution K, and an Applied Biosystems™ 7900HT Fast Real-Time PCR System at Institution F.

## Results

### Establishment of the one-day, single dose GHC prediction model

As shown in the Fig. 1, establishment of the GHC prediction model consisted of 5 steps: (1) Identifying 10 marker genes that were commonly altered in response to low doses of 5 GHCs using the microarray gene expression data from the Open TG-GATEs database. (2) Construction of the GHC predictive classifier using microarray expression data of marker genes from the low dose of the 3 GHCs, 2-AAF, NDEA, and 2-NF, as positive training data sets and microarray expression data of marker genes from low doses of the 14 OTHER compounds (listed in section 5) as negative training datasets. (3) In silico validation of the GHC predictive classifier. Microarray expression data of marker genes from the middle and high doses of the 3 GHCs used in step 2 (2-AAF, NDEA, and 2-NF) and microarray expression data from the low, middle, and high doses of 2 other GHCs, AFB1 and NMP, were used as GHC positive compounds. Microarray expression data of marker genes from the middle and high doses of the 14 OTHER compounds used in step 2 and low, middle, and high doses of 39 OTHER compounds were used as GHC negative compounds. (4) Determination of the usefulness of marker gene expression data measured by qPCR instead of microarray analysis (Validation Study 1): Evaluation of the marker gene expression of 5 GHCs by qPCR and microarray analysis and comparing the accuracy of prediction of the microarray analysis data and the qPCR data. (5) Multi-institutional validation: The predictive performance of the model was evaluated in multi-institutional one-day single oral administration studies (Validation Study 2). A total of 60 compounds including 19 GHCs were evaluated at 3 institutions. The validation used gene expression data of the marker genes, obtained through qPCR from these single oral administration studies.

**Fig. 1.**
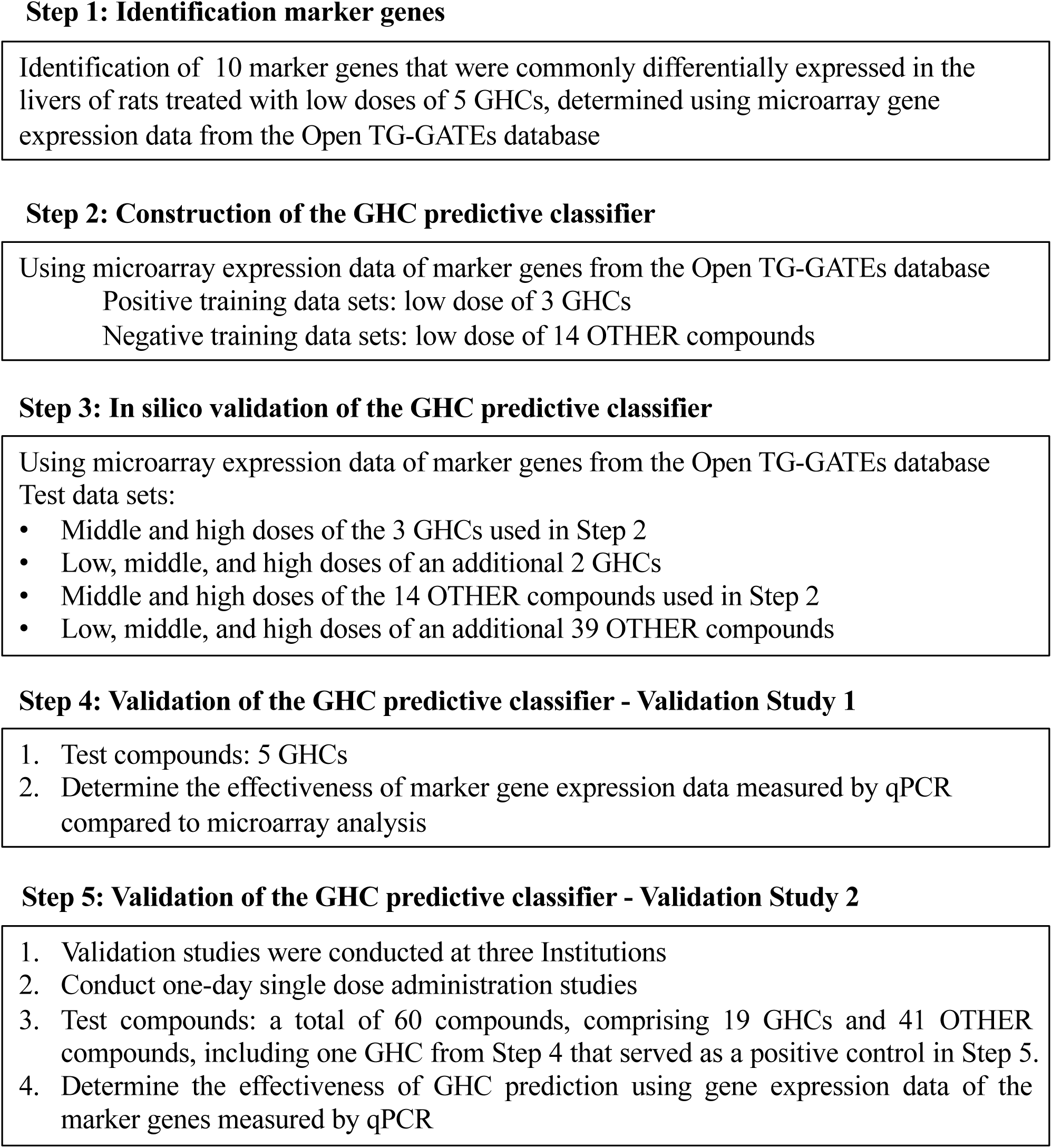
Workflow of the establishment of the one-day, single dose GHC prediction model.

### Identification of marker genes for GHC predictive classifier

As shown in the Fig. 2A, ten genes were identified, nine were upregulated and one was downregulated, that were commonly differentially expressed in the livers of rats treated with the 5 GHCs.

**Fig. 2.**
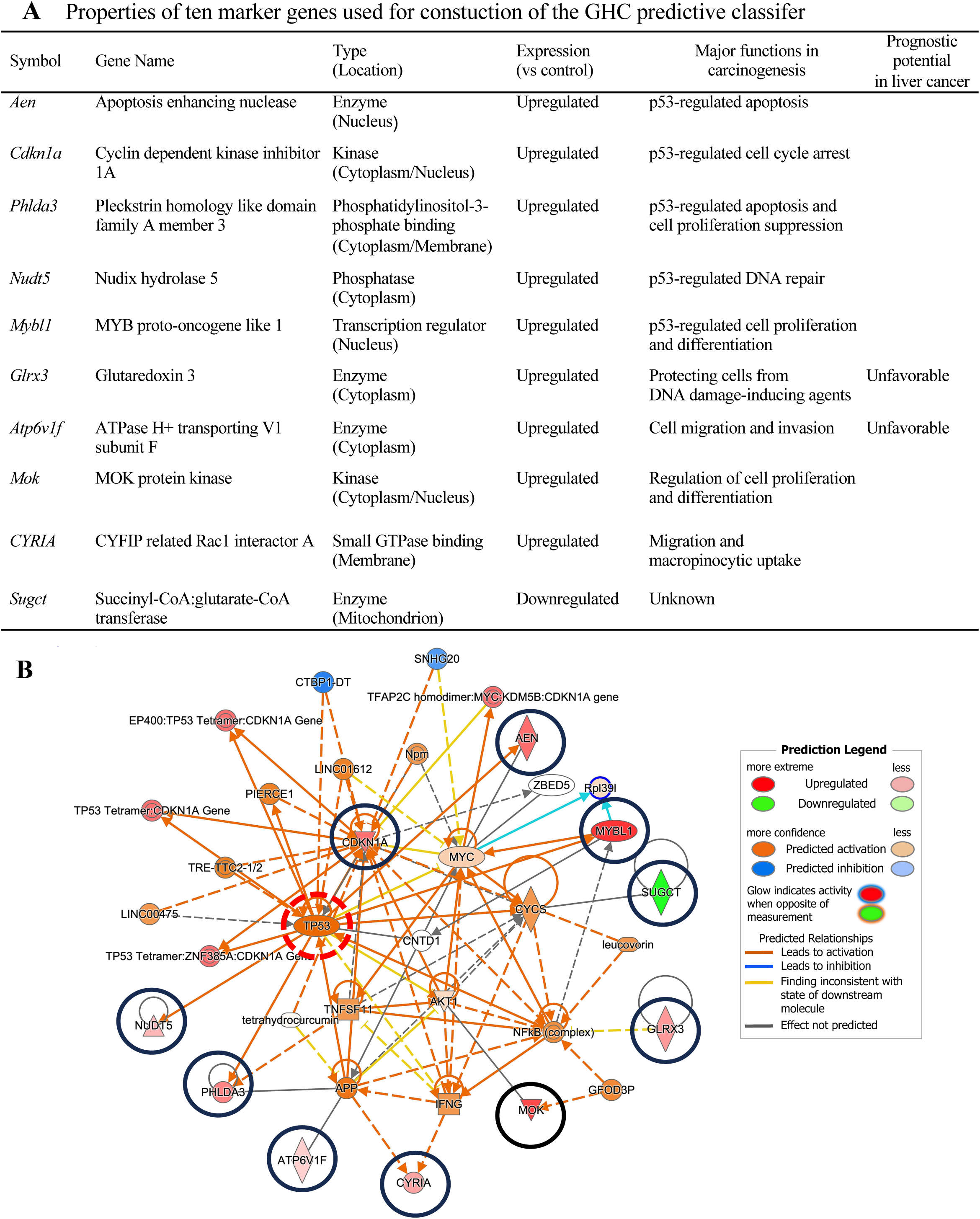
Properties and network analysis of marker genes used for GHC prediction in rat livers. (A) The properties of the ten marker genes used in construction of the GHC predictive classifier. (B) The network [Cancer, Cellular Growth and Proliferation, Organismal Injury and Abnormalities] generated by IPA analysis. The solid line circled genes represent the ten marker genes identified in the present study. An additional 25 molecules with predicted alternate expression are integrated into the network based on Ingenuity Pathways Knowledge Base.

To explore potential interactions among these ten genes, we conducted a network analysis using IPA software. These ten genes were considered focus genes and served as "seeds" for generating networks. As shown in Fig. 2B, IPA generated a single network labeled [Cancer, Cellular Growth and Proliferation, Organismal Injury and Abnormalities]. Notably, five of these ten genes, apoptosis enhancing nuclease (Vogel et al.), cyclin-dependent kinase inhibitor 1A (Cdkn1a), pleckstrin homology-like domain, family A, member 3 (Phlda3), nudix (nucleoside diphosphate linked moiety X)-type motif 5 (Nudt5), and myeloblastosis oncogene-like 1 (Mybl1), are regulated directly by the tumor suppressor protein p53 (Fig. 2B). The remaining five genes are MAPK/MAK/MRK overlapping kinase (Mok), glutaredoxin 3 (Glrx3), (succinyl-CoA: glutarate-CoA transferase (SUGCT), ATPase H+ transporting V1 subunit F (Atp6v1f), and CYFIP related Rac1 interactor A (CYRIA). The increased expression of the five p53-regulated genes one day after oral administration of the test compounds is considered a response to the genotoxic effects of GHCs, and alterations in the other five genes have been reported in various cancers, as will be discussed below.

In addition to the ten genes identified in this study, 25 molecules were integrated into the network from the QIAGEN Knowledge Base (Supplementary Table 3). The network [Cancer, Cellular Growth and Proliferation, Organismal Injury and Abnormalities] suggests that our ten marker genes do not act independently; instead, they interact with other cancer-related genes that may be GHC-specific. Therefore, we selected these ten genes as marker genes for constructing a classifier to predict GHCs.

### Construction of the GHC predictive classifier: Using Microarray Gene Expression Data from Training Set Compounds in the Open TG-GATEs

After using 3 GHCs and 14 OTHER compounds (listed in section 5) to train the GHC predictive classifier, the classifier was used to predict the genotoxic hepatic carcinogenicity of the training compounds. Low doses of the 3 GHCs were correctly predicted as positive, and all low doses of the 14 Other compounds in the training set were correctly predicted as negative. For the low doses of the training compounds, both sensitivity and specificity were 100% (3/3 and 14/14, respectively: these results are included in Table 2).

### In silico validation of the GHC classifier using microarray gene expression from the Open TG-GATEs

The results of the GHC predictive classifier, based on the training datasets, are shown in Table 2. The prediction results from the positive (low doses of 3 GHCs) and negative (low doses of 14 OTHER compounds) training sets were not included in the calculation of the sensitivity and specificity of the GHC predictive classifier. As indicated in Table 2, for the low doses of the tested compounds, both GHCs (AFB1 and NMP) were correctly classified as positive and all 39 OTHER compounds were correctly classified as negative, resulting in a sensitivity of 100% (2/2) and a specificity of 100% (39/39). For the middle doses of the tested compounds (5 GHCs and 53 OTHER compounds), both sensitivity and specificity were 100% (5/5 and 53/53, respectively). However, for the high doses of tested compounds (5 GHCs and 53 OTHER compounds), while the sensitivity remained at 100% (5/5), the specificity slightly decreased to 96% (51/53). The false positives at high doses were attributed to TAA, a NGHC, and MNU, a GNHC in rats. These results indicate the high sensitivity and specificity of the 10-gene-based GHC predictive classifier, across a dosing range from 1/10 of the MTD to the MTD, in single oral administration study.

### Validation Study 1: using 5 GHCs in one-day single dose oral administration studies

We assessed the effectiveness of the GHC predictive classifier using 5 GHCs not included in the Open TG-GATEs database. To reduce costs and analysis time, we also explored the potential of using qPCR as a replacement for microarray analysis in determining the expression data of marker genes.

Our analysis showed a significant correlation between the expression levels of ten marker genes measured by microarray and qPCR analyses at 1/3 of the LD50 dose (Supplementary Fig. 1). The prediction results for these 5 GHCs are summarized in Table 3. In the first experiment of Validation Study 1, four of the five GHC (2-Nitropropane (2-NP), 2,4-Diaminotoluene (2,4-DAT), MeIQx, and IQ) were correctly classified as positive at 1/3 of the LD50 dose using both microarray and qPCR data of the 10 marker genes. However, 2-AA was incorrectly predicted as negative at 1/3 of the LD50 dose using either microarray data or qPCR expression data of the 10 marker genes. Notably, in the second experiment, where only qPCR analysis was conducted, 2-AA was correctly classified as positive at the LD50 dose.

At 1/10 of the LD50 dose, three out of the five GHCs (2-NP, MeIQx, and IQ) were correctly classified as positive. However, at the lower dose of 1/30 of the LD50 dose none of the GHCs were classified as positive using qPCR data. The sensitivity was 0% (0/5), 60% (3/5), and 80% (4/5) at 1/33, 1/10, and 1/3 of the LD50 dose, respectively.

These findings indicate that qPCR analysis for marker gene expression is a viable alternative to microarray analysis. Furthermore, given that our GHC predictive classifier reached the highest sensitivity at 1/3 of the LD50 dose for the tested GHCs using qPCR expression data, this dose has been chosen as the starting point for subsequent validation studies. Doses will be escalated to the LD50 dose for any GHCs that are incorrectly classified as negative at lower doses.

### Validation Study 2: A multi-institutional validation study using qPCR gene expression data from 60 compounds in one-day single dose oral administration studies

To demonstrate the effectiveness and reliability of the GHC predictive classifier using qPCR data of 10 marker genes, we conducted an evaluation of a total of 60 compounds, comprising 19 GHCs and 41 OTHER compounds, in a series of one-day, single dose oral administration studies at three institutions - O, K, and F. Each independent study included groups treated with 2-NP and the vehicle solution as positive and negative controls, respectively. Based on the qPCR results of Validation Study 1 in which 2 out of 3 GHCs were correctly classified as positive when administered at 1/3 of the LD50 dose (as noted above, doses of MelQx and IQ were not determined by LD50), we adopted this dose as the initial dose for each compound in the initial series of studies. When 1/3 of the LD50 dose was more than 2000 mg/kg bw, the administered dose was 2000 mg/kg bw, the maximum limitation for acute oral toxicity studies according to the OECD Guidelines for the Testing of Chemicals (Test No. 420: Acute Oral Toxicity - Fixed Dose Procedure).

The sensitivity of the GHC predictive classifier for the 19 GHCs at 1/3 of the LD50 dose up to a maximum of 2000 mg/kg was 79% (15/19) (Table 4). Four GHCs were incorrectly classified as negative at this dose: DCA, VB, 4,4’-ODA, and HS. To determine whether the misclassifications of the 4 GHCs at 1/3 of the LD50 dose were due to the low dosage, we re-evaluated their carcinogenicity at higher doses. As summarized in Table 4, DCA was negative at 1/2 of the LD50 dose, but was correctly predicted as positive at 2/3 of the LD50 dose. HS was correctly classified as positive at both 1/2 and 2/3 of the LD50 doses. The marker gene expression for DCA and HS showed a dose-dependent increase between the 1/2 and 2/3 LD50 doses. VB and 4,4’-ODA were incorrectly classified as negative at both the 1/2 and 2/3 LD50 doses. Due to the lack of a dose-dependent response in marker gene expression at these doses for VB and 4,4’-ODA, we did not evaluate these compounds at higher doses.

The specificity of the GHC predictive classifier for the 41 OTHER compounds at 1/3 of the LD50 dose was 93% (38/41). Three OTHER compounds were incorrectly classified as positive at the 1/3 LD50 dose: MCT (a NGHC), ETH (a NGNC), and NHMI (a GNHC).

We verified the predictive results for three GNHCs (BZ, ETU, and HZ). These GNHCs were consistently classified as negative at doses of 1/3, 1/2, and 2/3 of the LD50 at 3 institutions, demonstrating high reproducibility of the GHC predictive classifier. The positive control, 2-NP, was reliably predicted as positive in all 11 independent studies conducted at the 3 institutions.

When considering multiple dosing levels, the combined sensitivity of the GHC predictive classifier for the 60 compounds in Validation study 2 increased to 89% (17/19), while the specificity remained stable at 93% (38/41) (Table 4). This increase suggests that sensitivity is enhanced when data from several doses are analyzed.

Furthermore, as summarized in Table 5, the combined sensitivity of the GHC predictive classifier for the 64 compounds—based on marker gene expression data obtained from qPCR analyses in Validation Studies 1 and 2—was 91% (21/23). The specificity was 93% (38/41), and the accuracy reached 92% (59/64). These results demonstrate the reliability and effectiveness of the GHC predictive classifier across a range of doses for various compounds.

**Table 5.** Summary of prediction results of the Validation studies 1 and 2 using marker gene expression data obtained by qPCR analysis.

|  | No. of<br>compounds tested | No of<br>compounds correctly classified |  |
| --- | --- | --- | --- |
| GHC <sup>a</sup> | 23 <sup>c</sup> | 21 | Sensitivity: 91% |
| OTHER <sup>b</sup> | 41 | 38 | Specificity: 93% |
| Total | 64 | 59 | Accuracy: 92% |

<sup>a</sup> GHC: genotoxic hepatocarcinogen.
<sup>b</sup> OTHER consists of NGHC: nongenotoxic hepatocarcinogens; NGNHC: nongenotoxic nonhepatocarcinogen (carcinogenic to organs other than the liver); NGNC: Nongenotoxic and noncarcinogen; GNHC; genotoxic nonhepatocarcinogen (carcinogenic to organs other than the liver); and GNC: genotoxic noncarcinogen.
<sup>c</sup> In Validation Study 1 five GHCs were evaluated and in Validation Study 2 nineteen GHCs were evaluated. As 2-NP was assessed in both studies (serving as a positive control in Validation Study 2), the total number of unique GHCs evaluated was 23.

### Comparison of prediction results between qPCR and microarray gene expression data

A total of 40 compounds, comprising 6 GHCs and 34 Other compounds, were evaluated using marker gene expression data from qPCR expression data obtained at Institutions O, K, and F (Table 4), microarray gene expression obtained at Institution O (Table 3: 2-NP, 2,4-DAT, MeIQx, IQ, and 2-AA), and microarray gene expression data obtained from Open TG-GATEs. The prediction results for all six GHCs (2-NP, 2,4-DAT, MeIQx, IQ, 2-AA, and NDEA) and 32 of 34 OTHER compounds obtained from qPCR gene expression data were consistent with the microarray gene expression data (Supplementary Table 4).

Among the 34 OTHER compounds, 32 showed consistent prediction results between qPCR and microarray analysis. However, discrepancies were observed for MCT and ETH, which were inaccurately predicted as positive using the qPCR data, whereas they were correctly classified as negative in the microarray analysis. This inconsistency was unexpected, especially since the doses used in the qPCR analysis were within the range of the doses used in the microarray analysis.

Overall, these results demonstrate that the expression data of the 10 marker genes derived from qPCR are highly consistent with microarray analysis data for 38 out of the 40 tested compounds, with 37 of these 38 compounds correctly classified, underlining the accuracy, efficacy, and robustness of the GHC predictive classifier when utilizing qPCR data.

## Discussion

Since exposure to chemical carcinogens is a significant risk factor for human cancers, reducing exposure to these substances can greatly contribute to cancer prevention. In the present study, we developed a novel support vector machine-based one-day, single-dose prediction model of GHCs in rats. We also demonstrated that expression data from 10 marker genes obtained via qPCR can be directly used for classification, replacing microarray analysis. Our model has shown high accuracy and reliability, with a sensitivity of 91% and specificity of 93% across a series of validation studies using 64 compounds in three institutions. Our prediction model is effective for rapid detection of GHCs, aiding in prioritizing compounds for further testing and informing strategies to reduce exposure to GHCs. Therefore, it has significant implications for predicting the safety of chemicals and pharmaceuticals.

The typical workflow of our one-day single oral dose prediction model is illustrated in Figure 3. Stage 1 (Two days for animal studies): On Day 1, administer a single oral dose of the test chemical to rats (5 rats per group). On Day 2, 24 hours after treatment, collect and store liver tissues for RNA analysis. Stage 2 (Two days for RNA extraction and qPCR analyses): Extract RNA from collected liver tissues and conduct RT-qPCR analyses. Stage 3 (One day for data analysis): Analyze qPCR expression data and perform prediction classification analysis. Our model offers several advantages, enabling high-throughput and high accuracy: (1) It is short-term, cost-effective, and labor-efficient. (2) Fewer rats are required, reducing animal use. (3) Only a small amount of the test chemical is needed. (4) qPCR-based gene expression analysis is convenient and less complex compared to microarray analysis, ensuring data reliability and reproducibility across laboratories. (5) It supports the 3R principles (Replacement, Reduction, and Refinement) in animal research.

**Fig. 3.**
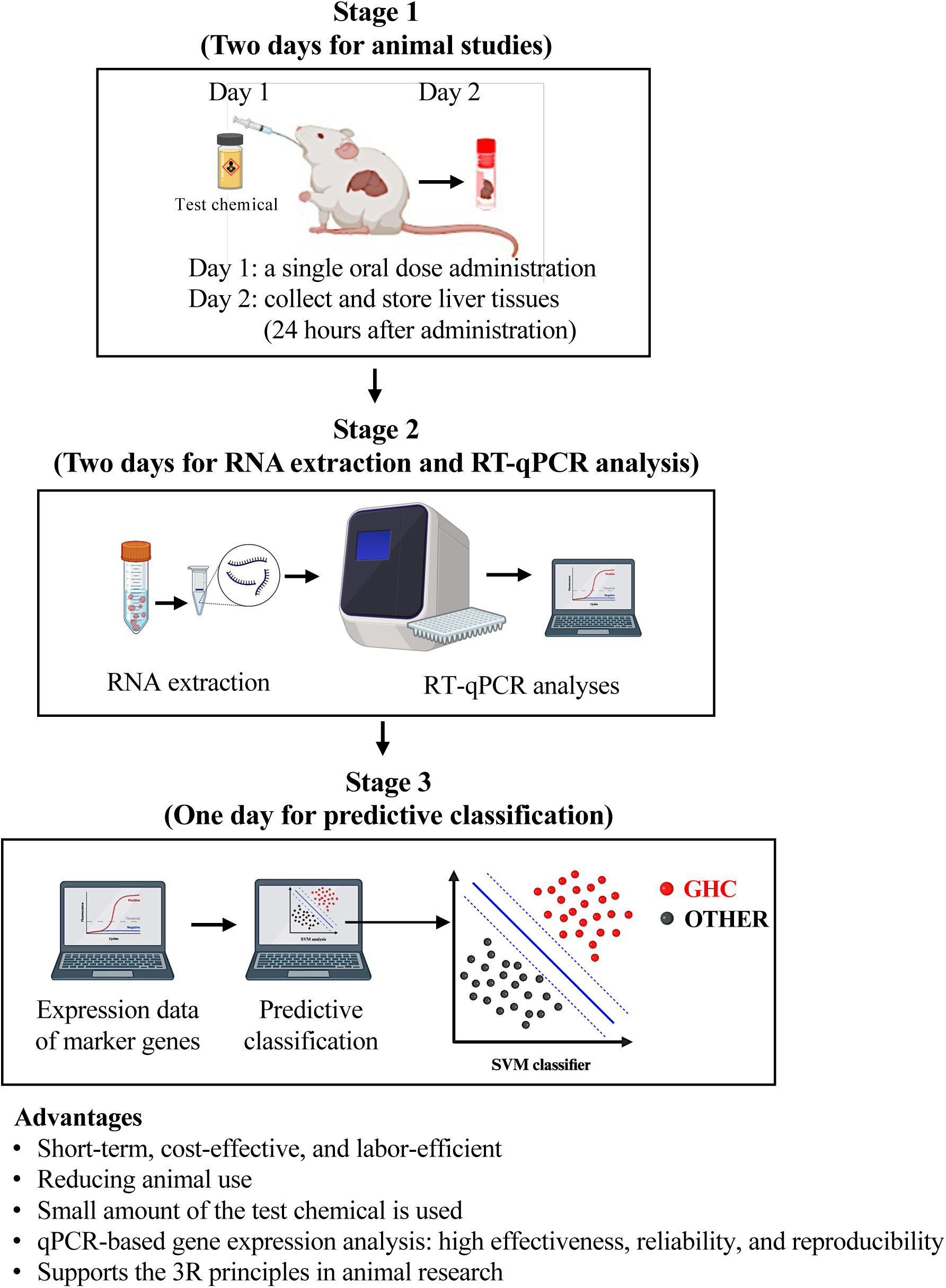
The typical workflow of one-day single dose prediction model (Created with BioRender.com.).

Regarding false negative predictions: 4,4’-ODA and VB, being GHCs in rats, were incorrectly classified as negative in our GHC prediction model (Table 4). Some possible explanations for this discrepancy include: (1) Limitations in marker gene selection: These compounds may induce carcinogenic pathways or gene expressions not covered by the current set of marker genes. (2) Route of exposure: The impact of the route of exposure on compound metabolism and genotoxic effects is significant. Inhalation exposure to VB, for instance, induced hepatocellular carcinoma and angiosarcoma in rat livers (Benya et al. 1982). Limited information on the carcinogenic effects of VB via oral administration suggests that its hepatocarcinogenicity might depend on the administration route. (3) Dosage: The sensitivity of our predictive model can be affected if the dosage used is below the effective dose needed to induce detectable changes in marker gene expression. However, 4,4’-ODA and VB were classified as negative at up to 1/3, 1/2, and 2/3 of their LD50 (Table 4), with no dose-response changes in marker gene expression, suggesting that our current set of marker genes may not be sufficiently sensitive to the specific genotoxic hepatocarcinogenic effects of these two compounds. Since the mechanism of action for GHCs is complex, a GHC might exert its carcinogenic effects through complex, multifactorial mechanisms that are not easily distinguished by a single predictive model. We are conducting ongoing studies to identify a complementary set of marker genes to refine the accuracy of GHC prediction.

Regarding the false positive prediction of NHMI: NHMI is genotoxic and induced squamous cell carcinoma (SCC) in the lungs of rats but was not carcinogenic to the liver (Reznik-Schüller and Gregg 1981). However, it was incorrectly classified as positive in the current study. Since the liver plays a crucial role in the metabolism of chemicals, gene expression alterations induced by chemicals can be complex and might not be accurately distinguished by a single prediction model. Utilizing a combination of complementary sets of marker genes could enhance the accuracy of prediction, as discussed above in the context of false negative predictions.

The cause of the discrepancy in the prediction results for MCT and ETH between the Open TG-GATEs microarray data and the qPCR expression data remains unclear. MCT, a non-genotoxic hepatocarcinogen, was predicted as negative at doses of 3, 10, and 30 mg/kg using microarray data from Open TG-GATEs (Table 2). However, in Validation Study 2 with qPCR mRNA expression data, MCT was predicted as positive at a dose of 20 mg/kg (Table 4). Similarly, ETH, which is non-genotoxic and non-carcinogenic, was predicted as negative at doses of 100, 300, and 1000 mg/kg using the Open TG-GATEs microarray data (Table 2), but was predicted as positive at a dose of 440 mg/kg in Validation Study 2 using qPCR mRNA expression data (Table 4). This inconsistency might be due to the differing sensitivities and specificities of qPCR and microarray techniques. However, it’s notable that the prediction results for 38 out of 40 compounds, which were classified using both Open TG-GATEs microarray data and our microarray and qPCR data, were consistent. Additional research is required to resolve these discrepancies and improve the accuracy of the predictive model.

As summarized in Figure 2 and described below, the 10 marker genes are aberrantly expressed in various human cancers. In particular, p53-regulated genes (Aen, Cdkn1a, Phlda3, Nudt5, and Mybl1) are aberrantly expressed in various human cancers, including liver cancers. Additionally, overexpression of two genes (Glrx3 and ATP6V1F) are involved in unfavorable prognosis in human liver cancers (Hu et al. 2023; Uhlen et al. 2017a; Uhlen et al. 2017b). This suggests that our prediction model is mechanism-based and relevant to human cancers. Such relevance enhances the potential value of our model in identifying hazards in humans.

Regarding the functions of the 10 marker genes: Aen is a downstream mediator of p53. It is implicated in the initiation of apoptosis following DNA damage (Kawase et al. 2008). Decreased expression of Aen has been observed in human liver cancers, suggesting its involvement in liver carcinogenesis (Hamza et al. 2015; Pinato et al. 2018). Cdkn1a is a p53-regulated tumor suppressor gene. It acts as a cyclin-dependent kinase inhibitor and plays a crucial role in p53-mediated cell cycle arrest in response to DNA damage (Engeland 2022; Kreis et al. 2019). Decreased Cdkn1a expression has been frequently observed in various cancers, including liver cancer (Hui et al. 1998; Ozturk et al. 2009). PHLDA3 is a p53-regulated tumor suppressor gene. It is known to promote apoptosis and suppress cell proliferation by negatively impacting the AKT signaling pathway, and its expression is induced by p53 in response to DNA damage (Chen and Ohki 2020). Loss of PHLDA3 is involved in various cancers (Chen and Ohki 2020; Saito et al. 2022). NUDT5 is a p53-regulated MutT-type DNA repair enzyme. It is important in cellular defense against oxidative DNA damage. It catalyzes the degradation of 8-oxoguanine, a DNA lesion resulting from oxidative modification of guanine, thereby inhibiting mutagenesis and maintaining genomic stability (Burnum et al. 2012; Hori et al. 2010; Ito et al. 2005). Mybl1 is a p53-regulated member of the MYB transcription factor family. It is essential for cell growth and differentiation (Rivlin et al. 2014; Yoon et al. 2002). Overexpression of Mybl1 has been linked to increased growth, angiogenesis, and metastasis in liver cancer (Xie et al. 2020; Zhu et al. 2022). Glrx3 is an oxidoreductase enzyme. It protects cells from DNA damage-inducing agents by activating ATR-dependent signaling pathways (Pandya et al. 2019). Overexpression of Glrx3 is observed in various human cancers (Li et al. 2018; Lu et al. 2013), including liver cancer (Mollbrink et al. 2014). Notably, overexpression of Glrx3 is associated with an unfavorable prognosis in liver cancer, suggesting its potential as a prognostic marker (Uhlen et al. 2017a). ATP6V1F is a component of the multisubunit enzyme V-ATPase. It is involved in acidifying eukaryotic intracellular organelles, a process critical for various cellular functions, including protein sorting, zymogen activation, and receptor-mediated endocytosis. Recent findings indicate that ATP6V1F contributes to the progression of liver cancer by promoting cell migration and invasion, and by inhibiting apoptosis in cancer cells (Hu et al. 2023). Notably, overexpression of ATP6V1F is associated with an unfavorable prognosis in liver cancer, suggesting its potential as a prognostic marker (Hu et al. 2023; Uhlen et al. 2017b). MOK belongs to the MAP kinase superfamily. It acts as a tumor suppressor gene in intestinal tumorigenesis (Chen et al. 2013). It is involved in the regulation of cell proliferation and differentiation of intestinal epithelial cells (Chen et al. 2013). CYRIA acts as a crucial RAC1-binding regulator, influencing the dynamics of lamellipodia and macropinocytic uptake (Machesky 2023). It plays an important role in moderating migration of cancer cells by controlling macropinosome formation and the regulation of integrin internalization (Le et al. 2021). SUGCT is a mitochondrial enzyme involved in synthesizing glutaryl-CoAis. It is the only gene among the ten marker genes that shows decreased expression in response to GHCs. Knockout of SUGCT results in diet-linked, age-related microbiome imbalances in mice, leading to a diabetes-like metabolic syndrome phenotype that includes hepatic steatosis (Niska-Blakie et al. 2020). However, the role of SUGCT in cancer remains unclear.

In conclusion, our study has successfully developed a novel, reliable, highly sensitive, and specific prediction model for assessing the genotoxic hepatocarcinogenicity of chemicals just 24 hours after a single dose oral administration of the test material. This model is anticipated to be a valuable tool in the early stages of cancer risk assessment, significantly enhancing the efficiency of identifying and screening potential GHCs. Further investigations involving a broader range of chemicals will aid in further validating the current model’s effectiveness. Moreover, integrating this GHC prediction model with other carcinogenicity prediction models could provide a comprehensive approach for assessing the carcinogenicity of compounds. Such a strategy not only increases the accuracy and reliability of carcinogenic assessments but also support ethical scientific practices by potentially reducing and refining the use of animals in experimental risk assessments.

## Acknowledgements

This work was supported by Health and Labour Sciences Research Grants from the Ministry of Health, Labour and Welfare of Japan (20KD0101 and 22KD1003), and a grant from Japan Society for the Promotion of Science (23K09652). Guiyu Qiu is supported by a scholarship from Ichikawa International Scholarship Foundation. Runjie Guo is supported by scholarships from Nishimura International Scholarship Foundation, Japan and Association for Promotion of Research on Risk Assessment, Japan. The authors gratefully acknowledge the technical assistance of Rie Onodera, Keiko Sakata, Yuko Hisabayashi, and Yukiko Iura (Department of Molecular Pathology, Graduate School of Medicine School, Osaka Metropolitan University Osaka, Japan).

---

**Supplementary Fig. 1.**
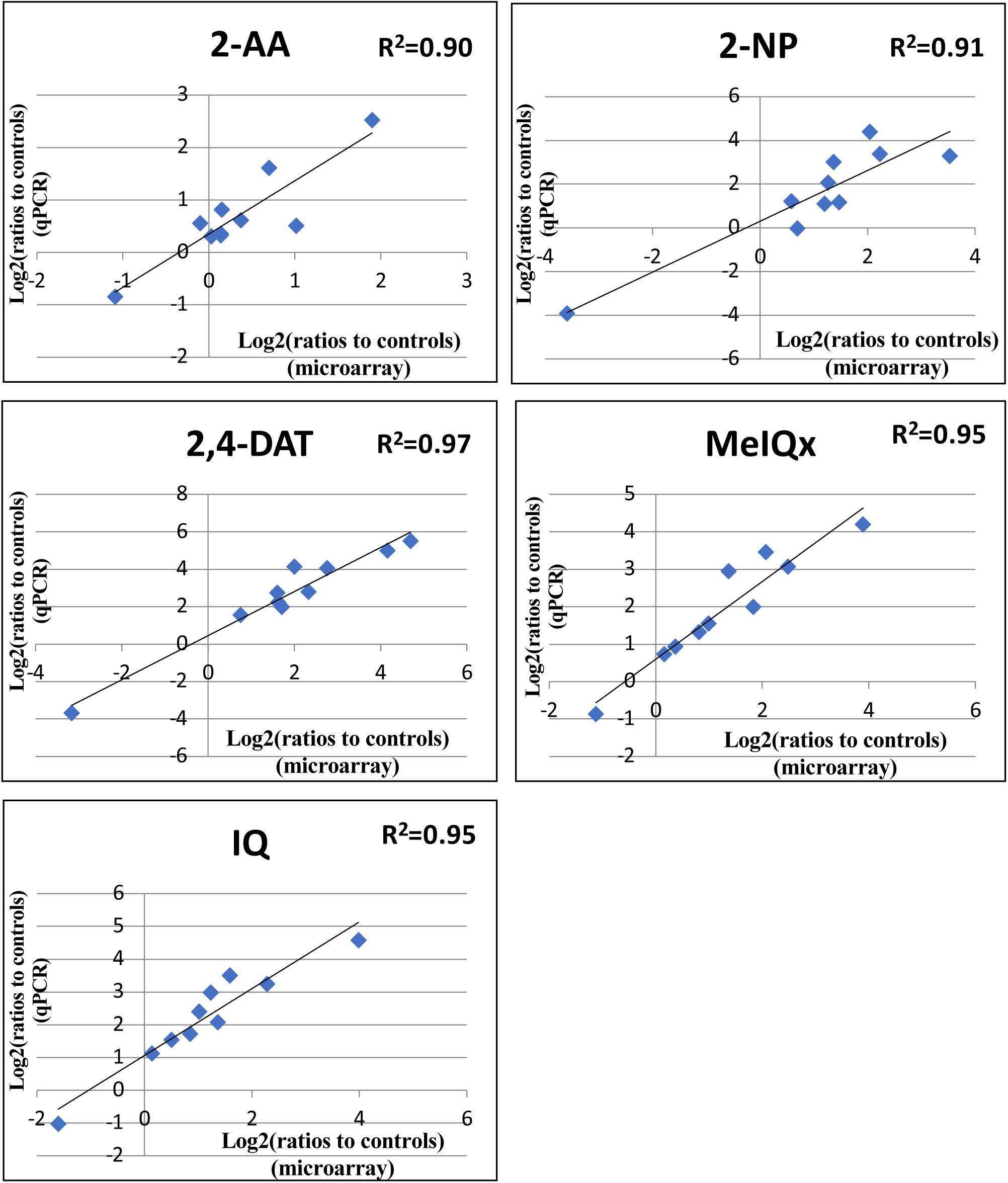
Relative expression levels of ten marker genes showed a significant correlation between microarray analysis and qPCR analysis of 5 GHCs administered at a dose of 1/3 of the LD50 dose. The X-axis represents the log2 (ratios to controls) values derived from microarray analysis; the Y-axis represents the log2 (ratios to controls) values from qPCR analysis. Each diamond-shaped point represents one of the ten marker genes.

**Supplementary Table 1.**
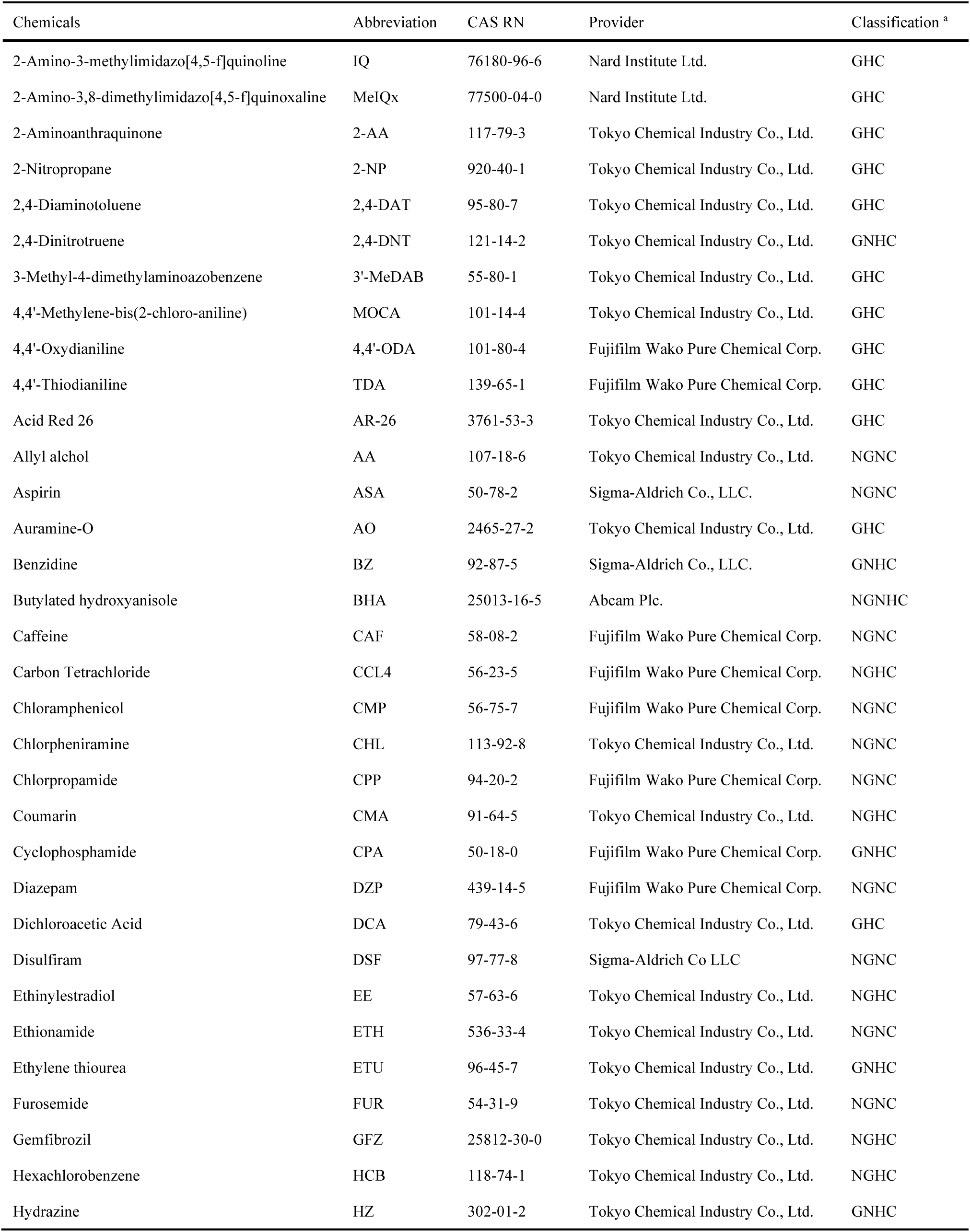
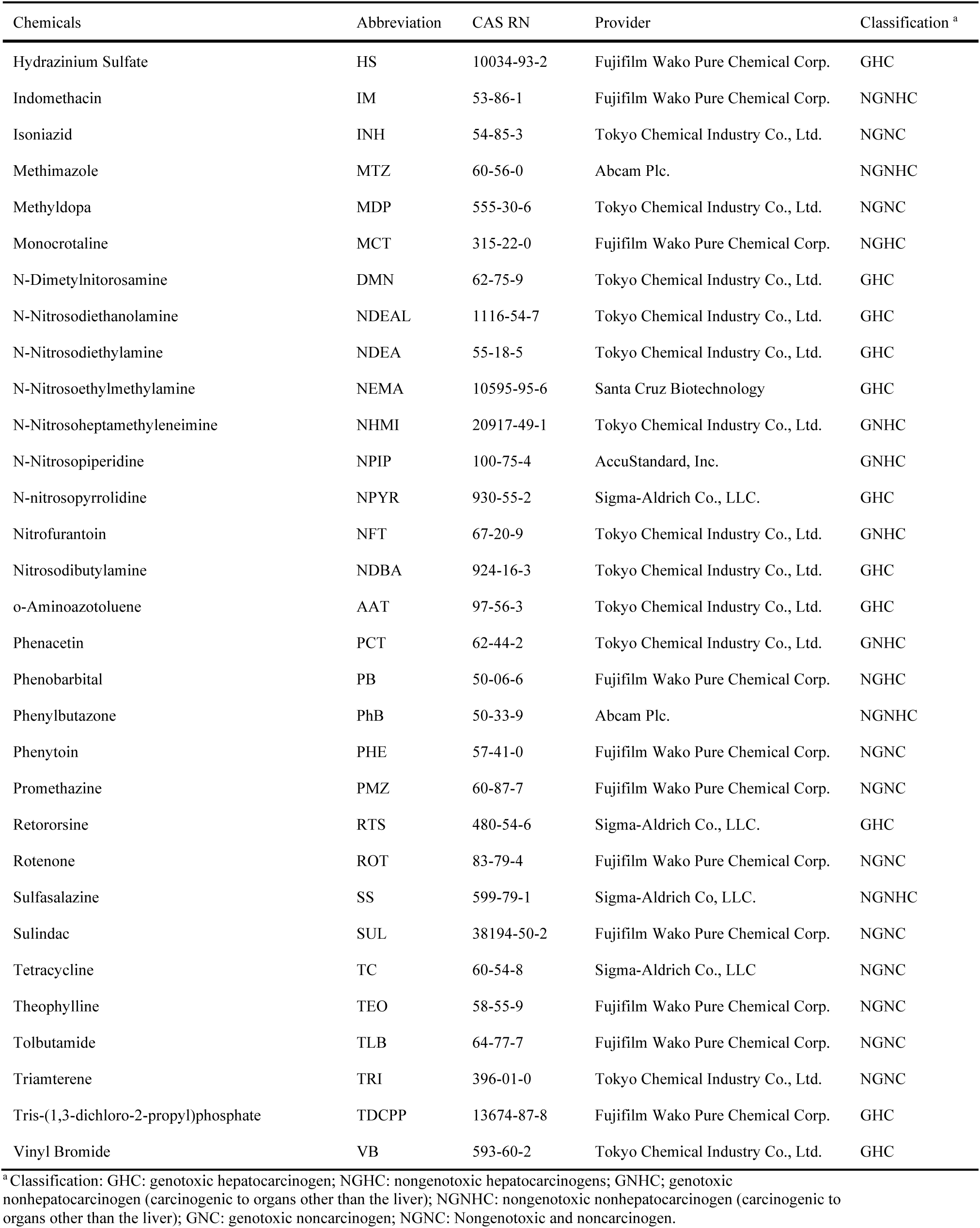
CAS numbers and suppliers of the chemicals tested in Validations Study 1 and Validation Study 2.

**Supplementary Table 2.**
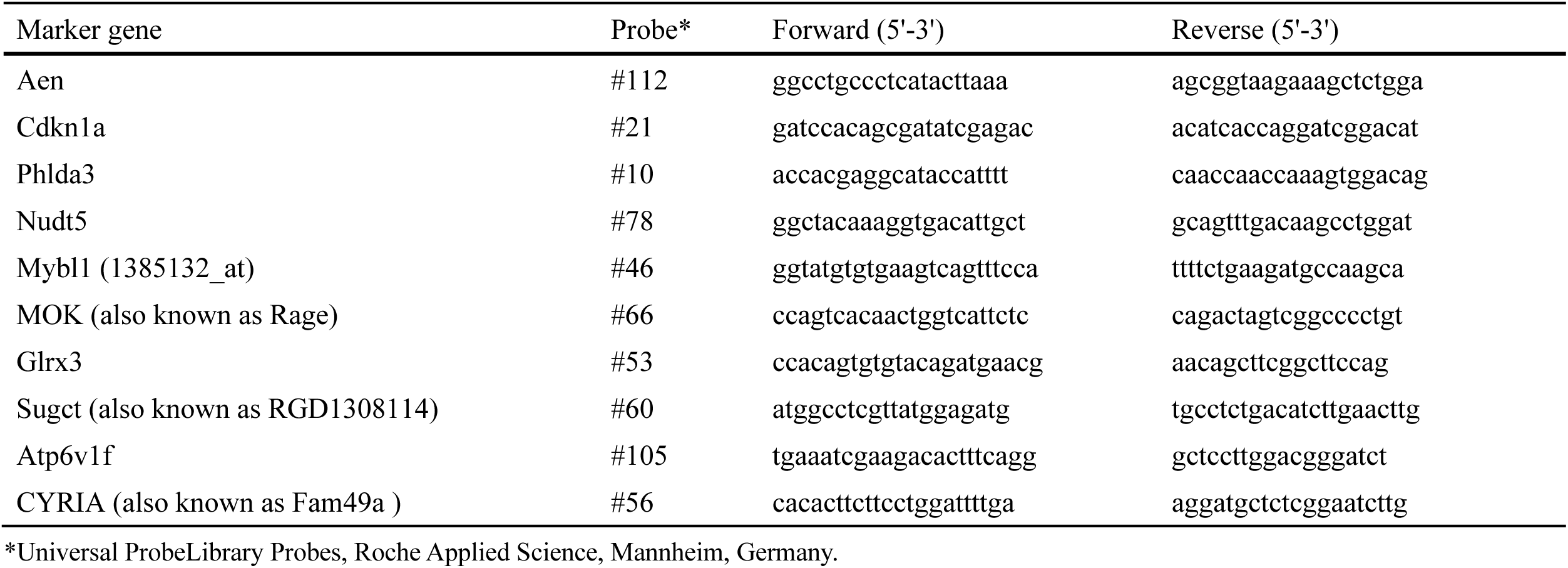
Primers and probes used in the qPCR.

**Supplementary Table 3.**
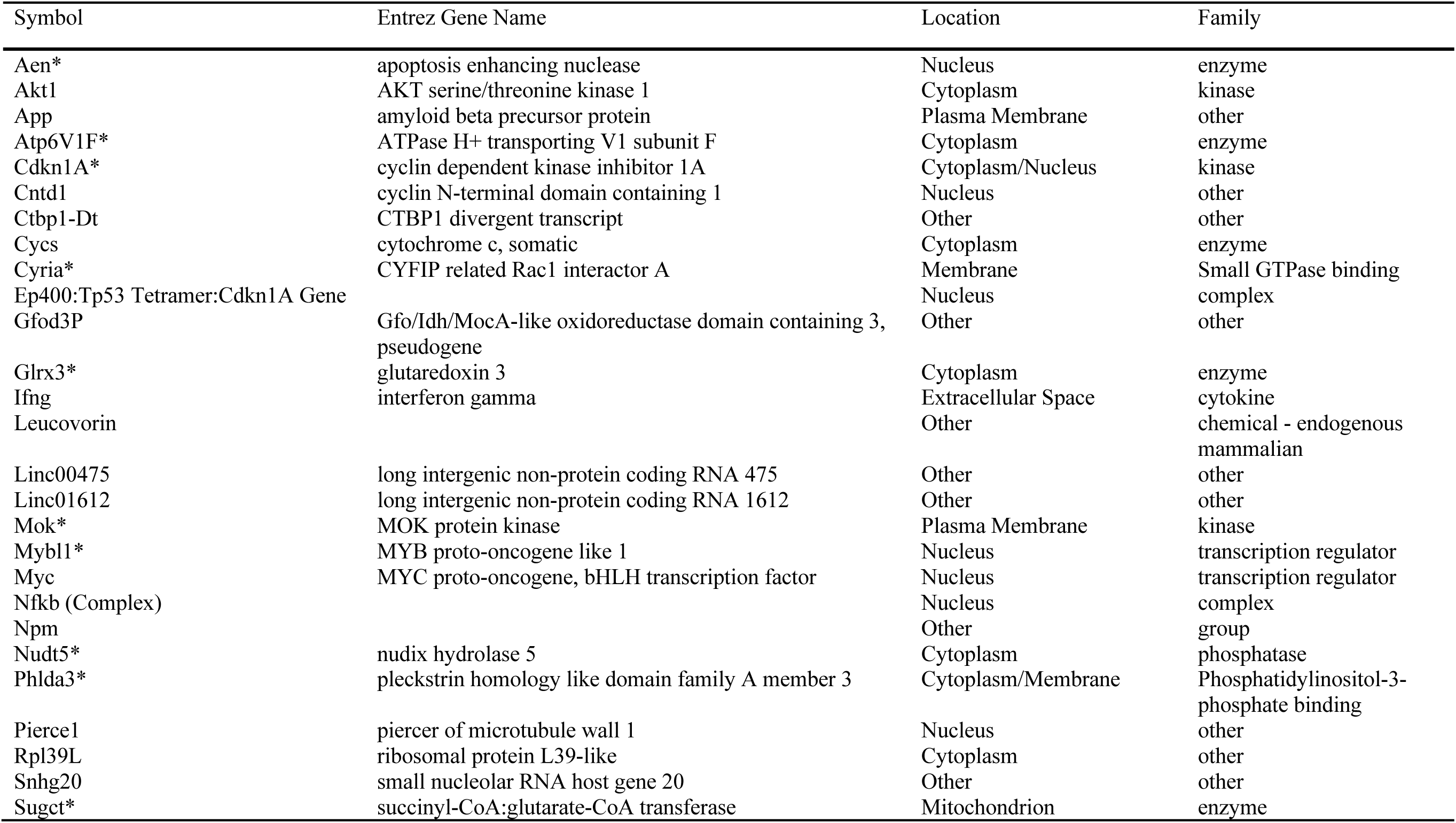
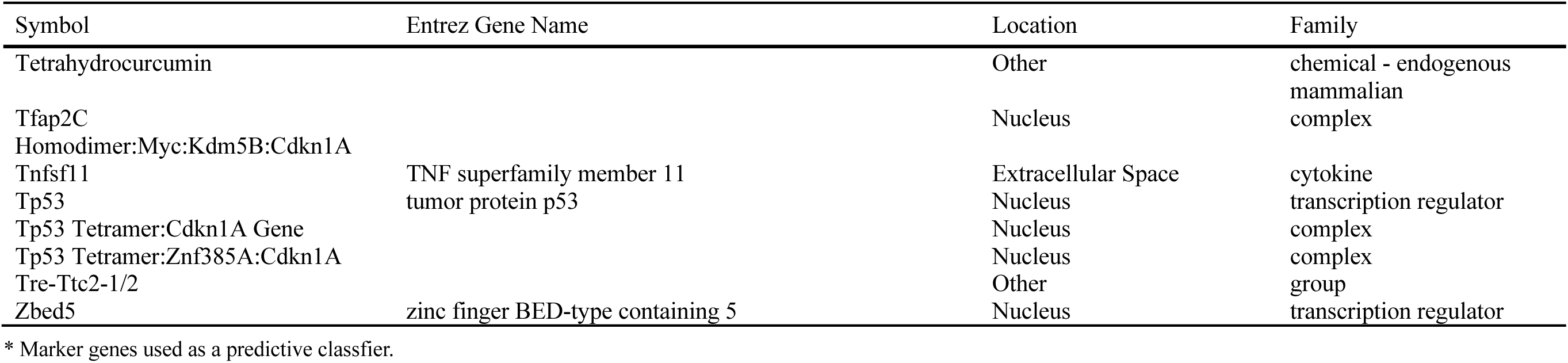
Molecules in the QIAGEN Knowledge Base network [Cancer, Cellular Growth and Proliferation, Organismal Injury and Abnormalities].

**Supplementary Table 4.**
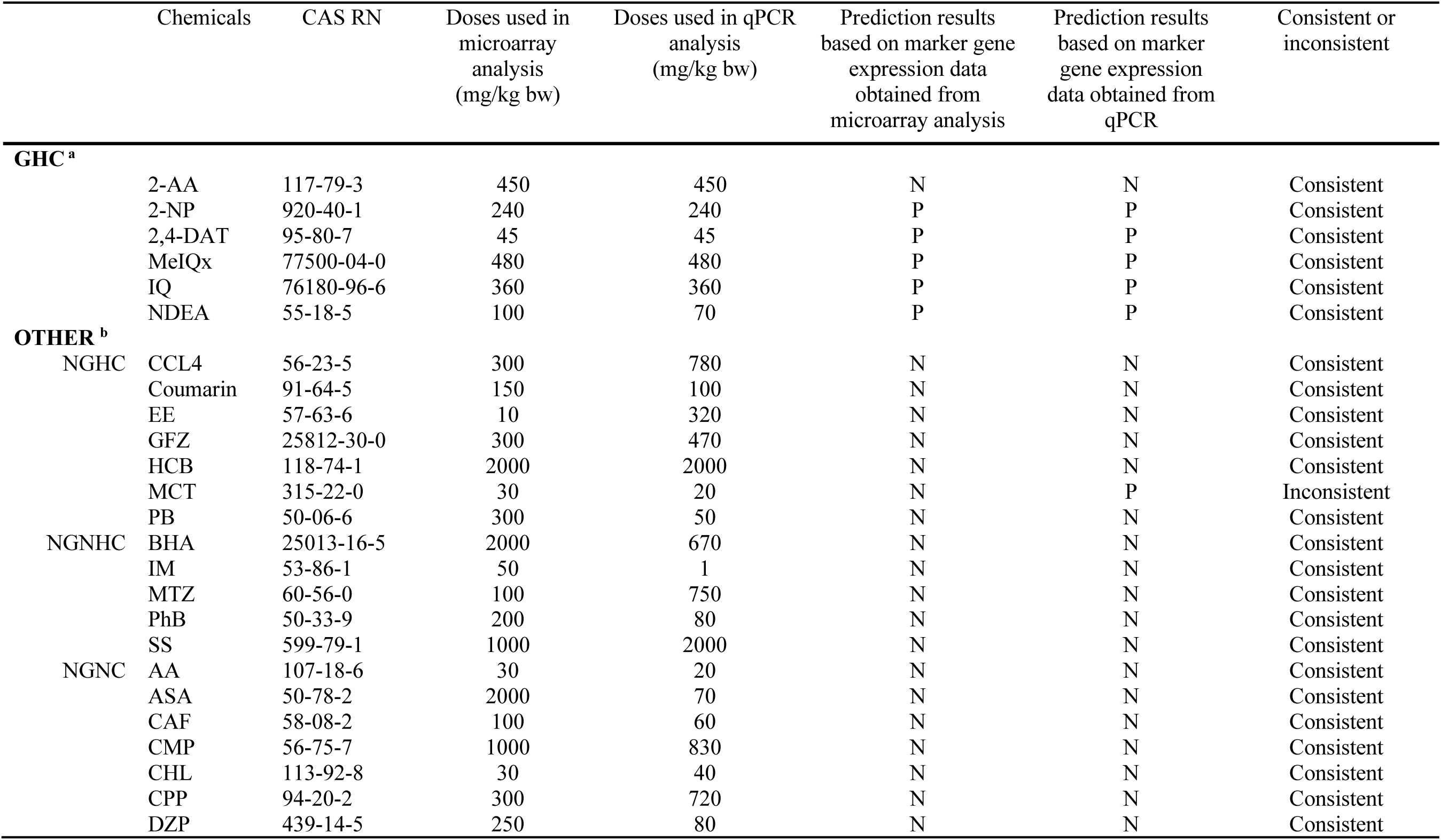
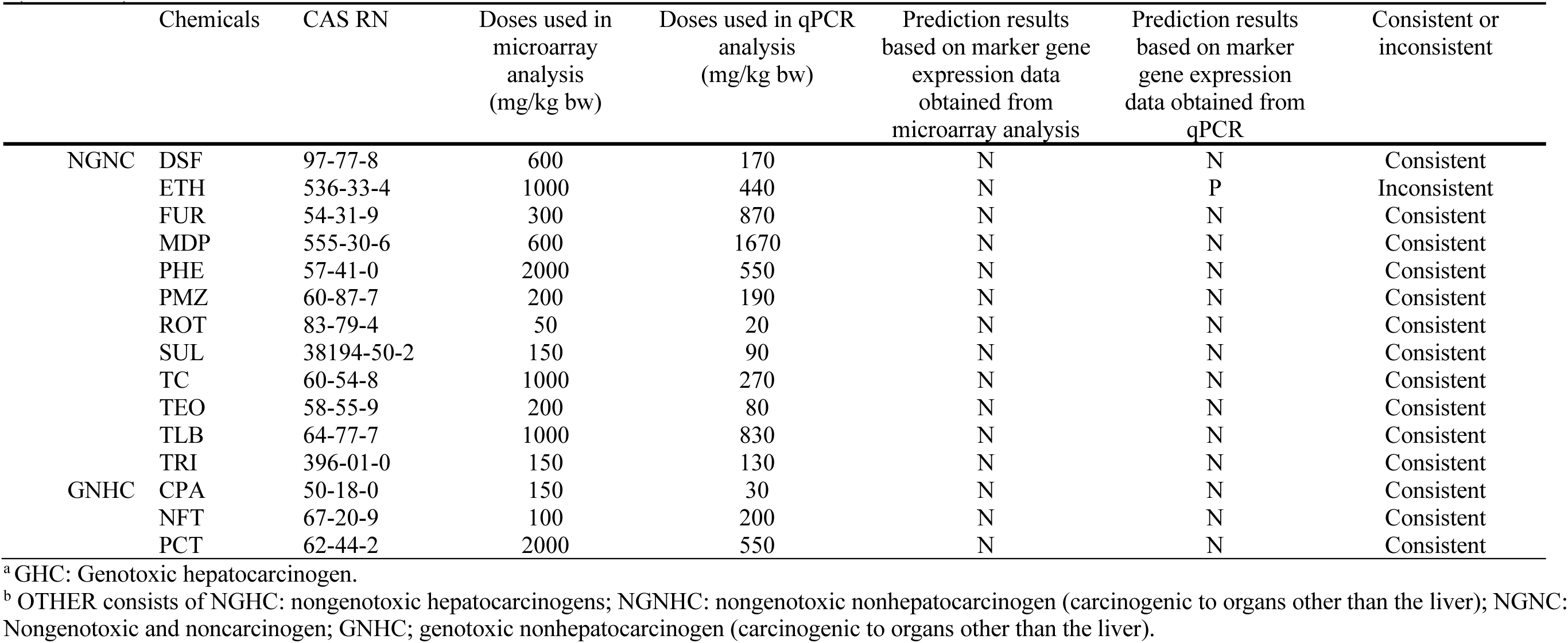
Comparison of prediction results using microarray gene expression data from Open TG-GATEs and qPCR gene expression data from Tables 3 and 4.

## Notes

### Competing Interest Statement

The authors have declared no competing interest.

## References

Ashby J (1996) Alternatives to the 2-species bioassay for the identification of potential human carcinogens. Hum Exp Toxicol 15(3):183–202 doi:10.1177/096032719601500301

Beal MA, Chen G, Dearfield KL, et al. (2023) Interpretation of In Vitro Concentration-Response Data for Risk Assessment and Regulatory Decision-making: Report from the 2022 IWGT Quantitative Analysis Expert Working Group Meeting. Environmental and molecular mutagenesis doi:10.1002/em.22582

Benya TJ, Busey WM, Dorato MA, Berteau PE (1982) Inhalation carcinogenicity bioassay of vinyl bromide in rats. Toxicology and applied pharmacology 64(3):367–79 doi:10.1016/0041-008x(82)90233-2

Bolt HM (2008) The Concept of “Practical Thresholds” in the Derivation of Occupational Exposure Limits for Carcinogens by the Scientific Committee on Occupational Exposure Limits (SCOEL) of the European Union. Genes and Environment 30(4):114–119 doi:10.3123/jemsge.30.114

Cohen SM (2010a) An enhanced 13-week bioassay: an alternative to the 2-year bioassay to screen for human carcinogenesis. Experimental and toxicologic pathology : official journal of the Gesellschaft fur Toxikologische Pathologie 62(5):497–502 doi:10.1016/j.etp.2009.06.011

Cohen SM (2010b) Evaluation of possible carcinogenic risk to humans based on liver tumors in rodent assays: the two-year bioassay is no longer necessary. Toxicologic pathology 38(3):487–501 doi:10.1177/0192623310363813

Cohen SM, Boobis AR, Dellarco VL, et al. (2019) Chemical carcinogenicity revisited 3: Risk assessment of carcinogenic potential based on the current state of knowledge of carcinogenesis in humans. Regul Toxicol Pharmacol 103:100–105 doi:10.1016/j.yrtph.2019.01.017

Cohen SM, Robinson D, MacDonald J (2001) Alternative models for carcinogenicity testing. Toxicol Sci 64(1):14–9 doi:10.1093/toxsci/64.1.14

Corton JC, Hill T, Sutherland JJ, Stevens JL, Rooney J (2020) A Set of Six Gene Expression Biomarkers Identify Rat Liver Tumorigens in Short-term Assays. Toxicol Sci 177(1):11–26 doi:10.1093/toxsci/kfaa101

Corton JC, Mitchell CA, Auerbach S, et al. (2022) A Collaborative Initiative to Establish Genomic Biomarkers for Assessing Tumorigenic Potential to Reduce Reliance on Conventional Rodent Carcinogenicity Studies. Toxicol Sci 188(1):4–16 doi:10.1093/toxsci/kfac041

Eastmond DA, Vulimiri SV, French JE, Sonawane B (2013) The use of genetically modified mice in cancer risk assessment: challenges and limitations. Crit Rev Toxicol 43(8):611–31 doi:10.3109/10408444.2013.822844

Ellinger-Ziegelbauer H, Aubrecht J, Kleinjans JC, Ahr HJ (2009) Application of toxicogenomics to study mechanisms of genotoxicity and carcinogenicity. Toxicology letters 186(1):36–44 doi:10.1016/j.toxlet.2008.08.017

Ellinger-Ziegelbauer H, Gmuender H, Bandenburg A, Ahr HJ (2008) Prediction of a carcinogenic potential of rat hepatocarcinogens using toxicogenomics analysis of short-term in vivo studies. Mutation research 637(1-2):23–39 doi:10.1016/j.mrfmmm.2007.06.010

Ennever FK, Lave LB (2003) Implications of the lack of accuracy of the lifetime rodent bioassay for predicting human carcinogenicity. Regul Toxicol Pharmacol 38(1):52–7 doi:10.1016/s0273-2300(03)00068-0

EPA (2005) United States Environmental Protection Agency’s guidelines for carcinogen risk assessment. https://www.epa.gov/risk/guidelines-carcinogen-risk-assessment.

Fielden MR, Adai A, Dunn RT, 2nd, et al. (2011) Development and evaluation of a genomic signature for the prediction and mechanistic assessment of nongenotoxic hepatocarcinogens in the rat. Toxicol Sci 124(1):54–74 doi:10.1093/toxsci/kfr202

Furihata C, Watanabe T, Suzuki T, Hamada S, Nakajima M (2016) Collaborative studies in toxicogenomics in rodent liver in JEMS.MMS; a useful application of principal component analysis on toxicogenomics. Genes and environment : the official journal of the Japanese Environmental Mutagen Society 38:15 doi:10.1186/s41021-016-0041-0

Gant TW, Auerbach SS, Von Bergen M, et al. (2023) Applying genomics in regulatory toxicology: a report of the ECETOC workshop on omics threshold on non-adversity. Archives of toxicology 97(8):2291–2302 doi:10.1007/s00204-023-03522-3

Gold LS, Manley NB, Slone TH, Ward JM (2001) Compendium of chemical carcinogens by target organ: results of chronic bioassays in rats, mice, hamsters, dogs, and monkeys. Toxicologic pathology 29(6):639–52 doi:10.1080/019262301753385979

Gold LS, Slone TH, Manley NB, Bernstein L (1991) Target organs in chronic bioassays of 533 chemical carcinogens. Environ Health Perspect 93:233–46 doi:10.1289/ehp.9193233

Hartwig A, Arand M, Epe B, et al. (2020) Mode of action-based risk assessment of genotoxic carcinogens. Archives of toxicology 94(6):1787–1877 doi:10.1007/s00204-020-02733-2

Hill T, Rooney J, Abedini J, El-Masri H, Wood CE, Corton JC (2020) Gene Expression Thresholds Derived From Short-term Exposures Identify Rat Liver Tumorigens. Toxicol Sci 177(1):41–59 doi:10.1093/toxsci/kfaa102

Hsieh JH, Smith-Roe SL, Huang R, et al. (2019) Identifying Compounds with Genotoxicity Potential Using Tox21 High-Throughput Screening Assays. Chemical research in toxicology 32(7):1384–1401 doi:10.1021/acs.chemrestox.9b00053

Hu X, Li D, Zhu H, Yu T, Xiong X, Xu X (2023) ATP6V1F is a novel prognostic biomarker and potential immunotherapy target for hepatocellular carcinoma. BMC Med Genomics 16(1):188 doi:10.1186/s12920-023-01624-6

ICH (2008) International Conference on Harmonisation. Guidance on Genotoxicity Testing and Data Interpretation for Pharmaceuticals Intended for Human Use (ICH Topic S2 (R1).

Ito N, Tamano S, Shirai T (2003) A medium-term rat liver bioassay for rapid in vivo detection of carcinogenic potential of chemicals. Cancer science 94(1):3–8 doi:10.1111/j.1349-7006.2003.tb01343.x

Kanki M, Gi M, Fujioka M, Wanibuchi H (2016) Detection of non-genotoxic hepatocarcinogens and prediction of their mechanism of action in rats using gene marker sets. The Journal of toxicological sciences 41(2):281–92 doi:10.2131/jts.41.281

Kirkland D, Aardema M, Henderson L, Müller L (2005) Evaluation of the ability of a battery of three in vitro genotoxicity tests to discriminate rodent carcinogens and non-carcinogens I. Sensitivity, specificity and relative predictivity. Mutation research 584(1-2):1–256 doi:10.1016/j.mrgentox.2005.02.004

Kirkland DJ, Aardema M, Banduhn N, et al. (2007) In vitro approaches to develop weight of evidence (WoE) and mode of action (MoA) discussions with positive in vitro genotoxicity results. Mutagenesis 22(3):161–75 doi:10.1093/mutage/gem006

Mateo EM, Tarazona A, Jimenez M, Mateo F (2022) Lactic Acid Bacteria as Potential Agents for Biocontrol of Aflatoxigenic and Ochratoxigenic Fungi. Toxins (Basel) 14(11) doi:10.3390/toxins14110807

Nohmi T (2018) Thresholds of Genotoxic and Non-Genotoxic Carcinogens. Toxicol Res 34(4):281–290 doi:10.5487/tr.2018.34.4.281

Nohmi T, Suzuki T, Masumura K (2000) Recent advances in the protocols of transgenic mouse mutation assays. Mutation research 455(1-2):191–215

NTP (2002) Final report on Carcinogens (RcC), Background Ducument for Selected Heterocyclic Amines: PhIP, MeIQ, and MeIQx. National Toxicology Program

O’Brien J, Renwick AG, Constable A, et al. (2006) Approaches to the risk assessment of genotoxic carcinogens in food: a critical appraisal. Food Chem Toxicol 44(10):1613–35 doi:10.1016/j.fct.2006.07.004

Reznik-Schüller HM, Gregg M (1981) Pathogenesis of lung tumors induced by N-nitrosoheptamethyleneimine in F344 rats. Virchows Archiv A 393:333–343

SCHER (2009) Risk assessment methodologies and approaches for genotoxic and carcinogenic substances (European Commission, Scientific Committee on Health and Environmental Risks) http://ec.europa.eu/health/ph_risk/committees/04_scher/docs/scher_o_113.pdf.

Smith BP, Auvil LS, Welge M, et al. (2020) Identification of early liver toxicity gene biomarkers using comparative supervised machine learning. Scientific reports 10(1):19128 doi:10.1038/s41598-020-76129-8

Takayama S, Nakatsuru Y, Masuda M, Ohgaki H, Sato S, Sugimura T (1984) Demonstration of carcinogenicity in F344 rats of 2-amino-3-methyl-imidazo[4,5-f]quinoline from broiled sardine, fried beef and beef extract. Gan 75(6):467–70

Uehara T, Minowa Y, Morikawa Y, et al. (2011) Prediction model of potential hepatocarcinogenicity of rat hepatocarcinogens using a large-scale toxicogenomics database. Toxicology and applied pharmacology 255(3):297–306 doi:10.1016/j.taap.2011.07.001

Uhlen M, Zhang C, Lee S, et al. (2017a) A pathology atlas of the human cancer transcriptome Science 357(6352) doi:10.1126/science.aan2507. Data available from v23.0.proteinatlas.org. (https://www.proteinatlas.org/ENSG00000108010-GLRX3/pathology/liver+cancer)

Uhlen M, Zhang C, Lee S, et al. (2017b) A pathology atlas of the human cancer transcriptome. Science 357(6352) doi:10.1126/science.aan2507. Data available from v23.0.proteinatlas.org. (https://www.proteinatlas.org/ENSG00000128524-ATP6V1F/pathology/liver+cancer)

Vogel EW, Nivard MJ, Ballering LA, et al. (1996) DNA damage and repair in mutagenesis and carcinogenesis: implications of structure-activity relationships for cross-species extrapolation. Mutation research 353(1-2):177–218 doi:10.1016/0027-5107(96)00032-2

Waters MD, Jackson M, Lea I (2010) Characterizing and predicting carcinogenicity and mode of action using conventional and toxicogenomics methods. Mutation research 705(3):184–200 doi:10.1016/j.mrrev.2010.04.005

WHO, FAO (2021) Evaluation of certain food additives: eighty-ninth report of the Joint FAO/WHO Expert Committee on Food Additives. Geneva: World Health Organization and Food and Agriculture Organization of the United Nations. WHO; FAO;

Yamada F, Sumida K, Saito K (2016) An improved model of predicting hepatocarcinogenic potential in rats by using gene expression data. J Appl Toxicol 36(2):296–308 doi:10.1002/jat.3184

